# Estimating Muscle Parameters via Hierarchical Bayesian Neuromechanics

**DOI:** 10.64898/2026.07.08.737322

**Authors:** Russell T. Johnson, Yi Yu, Yannick Darmon, Victor R. Barradas, Dongze Ye, Nicolas Schweighofer, James M. Finley

## Abstract

Musculoskeletal models are widely used to relate muscle mechanics to movement patterns in biomechanics. Accurate estimation of muscle parameters is essential for building individualized models, yet most rely on generic parameters derived from cadaveric data that do not reflect subject-specific properties critical to force generation. Here, we introduce a hierarchical Bayesian framework that leverages surface electromyography (EMG) and torque data from isometric elbow tasks to estimate subject-specific muscle parameters, overcoming limitations of generic parameter sets. This approach accounts for both inter-individual variability and uncertainty in measurement and model structure. The model infers six key parameters per subject, including flexor and extensor muscle strength, tendon slack length, moment arm geometry, and nonlinear EMG-to-activation relationships.

We estimated model parameters for 14 young, healthy adults performing isometric elbow flexion and extension at multiple joint angles and torque levels. The six-parameter hierarchical-Bayesian musculoskeletal model accurately reproduced measured net elbow torque (R² = 0.96) and outperformed simpler configurations. Muscle strength parameters varied substantially across individuals, from approximately 1.0 to 3.5. On average, participants exhibited about twice those of the OpenSim 26 generic model. In contrast, tendon slack length estimates varied minimally across subjects. Bilateral testing revealed moderate correlations between left- and right-arm parameters, supporting the model’s ability to capture subject-specific anatomical features. Cross-validation confirmed robust predictive performance, and convergence diagnostics indicated reliable sampling.

Compared to traditional EMG-driven or imaging-based personalization methods, our approach quantifies uncertainty, enables partial pooling across subjects, and avoids reliance on invasive or time-intensive measurements. The framework is extensible to dynamic tasks and adaptable to clinical populations, including individuals post-stroke. These results demonstrate that hierarchical Bayesian inference can robustly personalize musculoskeletal models and advance our understanding of biomechanics.

## 1. Background

Understanding how muscle forces are generated and distributed during human movement is a central challenge in biomechanics and motor control. Musculoskeletal models provide a powerful framework for estimating muscle forces from experimental data, but these estimates are highly sensitive to model parameters and assumptions (Bernstein, 1967; Erdemir et al., 2007; Simpson et al., 2015; Stanev & Moustakas, 2019; Valente et al., 2014). Parameters such as peak isometric force and tendon stiffness are often derived from cadaveric data, which may not reflect the physiology of the individuals being studied, especially in young, healthy populations (Ward et al., 2009).

To address this limitation, researchers have developed personalized musculoskeletal models using various imaging techniques such as MRI or ultrasound (Blemker et al., 2007; Li & Tong, 2016; Valente et al., 2017). MRI-based musculoskeletal models, which personalize parameters such as bone geometry, muscle paths, and muscle architecture, have demonstrated improved anatomical fidelity and force estimation compared to scaled generic models (Blemker et al., 2007; Correa et al., 2011; Modenese & Kohout, 2020; Scheys et al., 2011), but they are expensive and often require extensive imaging and segmentation workflows, limiting their scalability in experimental and clinical settings.

An alternative strategy is to infer subject-specific muscle parameters directly from experimental data using statistical modeling. Electromyography (EMG)-driven musculoskeletal models offer one such approach. Surface EMG reflects the electrical activity of muscle fibers during activation, arising from both descending central motor commands and reflex pathways driven by sensory feedback. EMG-driven musculoskeletal models use measured muscle electrical activity to adjust model parameters (e.g., tendon slack length, flexor and extensor strength) while estimating muscle forces and joint moments (Lloyd & Besier, 2003; Shin et al., 2009). Foundational work by Lloyd and colleagues demonstrated that EMG signals can be used as an input to Hill-type muscle models and can predict joint kinetics with high accuracy (Buchanan et al., 2005; Pizzolato et al., 2015; Sartori et al., 2012). A common limitation across both MRI-based and EMG-driven personalization approaches, however, is that they typically yield single-point estimates of model parameters, often derived via maximum likelihood optimization, without quantifying the uncertainty associated with those estimates. This lack of uncertainty characterization can undermine confidence in the inferred parameters and limit the interpretability of model predictions, especially in clinical or experimental decision-making.

Bayesian inference offers a framework for estimating model parameters while accounting for uncertainty in both measurements and model structure (van Ravenzwaaij et al., 2018). Rather than producing a single point estimate of model parameters, Bayesian methods generate probability distributions over possible parameter values, reflecting both prior knowledge about a parameter and the evidence provided by the data (Andrew & Donald, 1992; Gelman et al., 2013). This approach allows researchers to quantify uncertainty in parameter estimates. In complex models such as Hill-type models, Markov Chain Monte Carlo (MCMC) methods, which iteratively generate parameter proposals and evaluate their likelihood given the data and priors (Andrieu et al., 2003; Brooks, 2011), are used to sample from these distributions. Over many iterations, MCMC builds a representative sample from the posterior distribution, enabling robust inference even in high-dimensional or non-linear models. This makes Bayesian inference particularly well-suited for musculoskeletal modeling, where uncertainty in experimental measurements, model structure, and physiological variability can affect outcomes such as joint kinematics, muscle forces, and joint contact forces (Ackland et al., 2012; Hicks et al., 2015; Myers et al., 2015; Raikova & Prilutsky, 2001; Scovil & Ronsky, 2006; Zuk et al., 2018). In previous work, we demonstrated the feasibility of using Bayesian methods to estimate plausible muscle forces that reproduce observed kinematics in a simulated elbow flexion-extension task (Johnson et al., 2022). However, that approach focused on the uncertainty in the objective function used to distribute muscle forces across multiple muscles (Crowninshield & Brand, 1981), rather than on the model parameters defining underlying muscle properties.

Several key parameters in musculoskeletal models—peak isometric force, tendon slack length, moment arm geometry, and the nonlinear relationship between EMG and muscle activation—vary substantially across individuals and have important implications for muscle function and force estimation. Peak isometric force varies with individual differences in muscle architecture, age, and muscular training history (Buchanan, 1995; Erskine et al., 2010; Merrigan et al., 2018; Nogueira et al., 2013). Tendon slack length can also differ between individuals and influence muscle-tendon dynamics and force transmission (Manal & Buchanan, 2004). Indeed, both peak isometric force and tendon slack length influence estimated muscle forces, muscle induced accelerations, and joint contact forces in musculoskeletal modeling simulations (Ackland et al., 2012; Myers et al., 2015; Winby et al., 2009). Similarly, muscle moment arms vary with joint angle and anthropometry (Murray et al., 2002), which affects torque generation. Finally, the relationship between surface EMG and muscle activation is nonlinear, subject-specific, and varies across muscles, particularly at low activation levels (Lloyd & Besier, 2003; Manal & Buchanan, 2003). Together, these findings support the need for individualized estimates for these parameters in biomechanical models, particularly when aiming to capture physiologically meaningful variability across a population.

In this study, we use Bayesian inference to estimate subject-specific muscle model parameters from EMG and torque data during isometric elbow contractions. We used a hierarchical Bayesian model that incorporates estimates of parameter uncertainty at both the individual-level and group-level, enabling robust parameter estimation across our experimental population (Gelman et al., 2013; McGlothlin & Viele, 2018; Schweighofer et al., 2023). We estimated model parameters to reproduce the measured elbow torque and EMG data from a set of healthy participants. We then validated our approach using synthetic data with known ground-truth parameters.

## 2. Methods

### 2.1 Human Subject Testing

A total of 15 healthy young individuals were recruited and provided written informed consent to participate in the experiment approved by the USC Institutional Review Board. All participants were tested on their dominant (right) side, and 8 of these participants were also tested on their left side. One participant was excluded from further analysis due to a lack of detectable biceps muscle modulation, likely caused by faulty EMG recording. Consequently, data from 14 participants (7 females; age = 28 ± 1 years; height = 1.69 ± 0.10 m; mass = 69.9 ± 11.6 kg) were included in the analysis. Anthropometric data were collected and used to scale the parameters of a musculoskeletal model.

Participants were seated in a chair with their trunk secured by a four-point seat belt and positioned close to a custom arm apparatus that restricted movement to elbow flexion and extension (Figure 1A and B). The forearm was strapped in a neutral pronation/supination position, and the position of the handle (hand grip) was adjusted such that the axis of rotation of the apparatus aligned with the elbow’s joint axis. The height and length of the apparatus were adjusted so the hand was ∼5 cm below the acromion and the shoulder was at ∼60° horizontal adduction.

**Figure 1:**
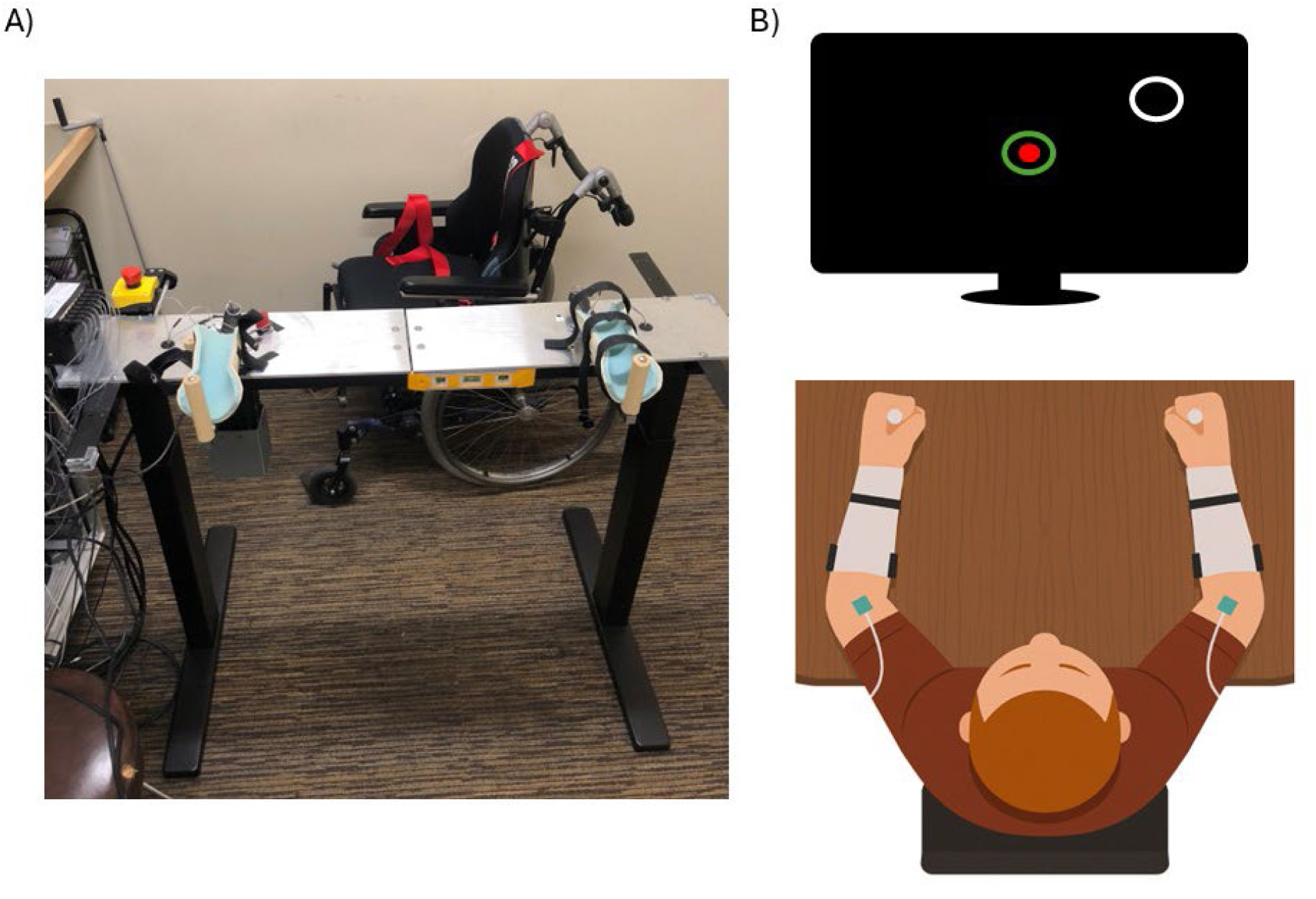
A) A photograph of the table and arm support set up used in the experimental data collection. B) A schematic of the experimental set up, with a participant in the arm supports, grasping the handles on each support. EMG electrodes are attached to relevant muscles. The participant views a television screen which displays feedback about the target force magnitude and direction.

Surface EMG was recorded from five elbow muscles: Biceps Brachii (flexor), Brachioradialis (flexor), Brachialis (flexor) (Date et al., 2021; Staudenmann & Taube, 2015), Triceps Brachii long head (extensor), and Triceps Brachii lateral head (extensor) using active bipolar electrodes (Biometrics Ltd, UK). EMG signals were acquired by the Biometrics system at 1 kHz and output as analog signals to a data acquisition device (USB-6259, National Instruments). The EMG signals were digitized synchronously at 2 kHz, along with position and force data.

The forearm was positioned at 20°, 70°, or 120° of elbow flexion, with the testing order randomized across participants. Elbow torque was measured using a calibrated one-axis force cell. For each posture, participants performed a maximum voluntary force (MVF) assessment followed by two blocks of submaximal trials, separated by 3 min rest periods. During MVF assessment, participants were instructed to perform two maximum isometric flexion and extension trials (∼5 s each). Separate MVF references for flexion and extension directions were defined as the mean of the 95th percentile force across trials for each direction. Following MVF assessment, participants performed sub-maximal isometric trials at 10%, 25%, and 40% of MVF in both flexion and extension directions. Each trial consisted of moving a cursor from the center of the screen to a target, stabilizing the cursor inside the target for 1 s, holding it for an extra 1 s, then returning to the center. This task was adapted from previously published isometric reaching paradigms (Barradas et al., 2020). Each participant completed 180 submaximal trials (3 elbow postures × 3 force magnitudes × 2 force directions × 5 trials × 2 blocks) and 12 MVF trials (3 elbow postures × 2 force directions × 2 trials).

EMG signals were band-pass filtered (20–350 Hz, 2nd-order, zero-phase forward–backward Butterworth filter), rectified, and low-pass filtered at 5 Hz (2nd-order, zero-phase forward–backward Butterworth filter) to obtain EMG envelopes. Force signals were low-pass filtered at 20 Hz (2nd-order, zero-phase forward–backward Butterworth filter). For each submaximal trial, EMG and force data were analyzed during the target hold period (final 1 s of the hold phase). For consistency, MVF trials were analyzed using a 1-s window centered around peak force. For each posture and direction (flexion and extension), only the MVF trial with the highest force was retained for analysis. EMG normalization was performed using the greatest mean EMG value observed across MVF trials. The mean and standard deviation of force and normalized EMG were computed within the window of interest for each trial and used in subsequent modeling.

### 2.2 Musculoskeletal Model

Muscle forces were modeled using a Hill-type formulation consisting of a contractile element (CE) and a series elastic element (SEE). In the CE, force is determined by three components: the maximal isometric force, the force–length relationship, and muscle activation. In the SEE, force is determined by tendon properties, including tendon slack length and stiffness. Activation is obtained from EMG via a nonlinear EMG-to-activation mapping.

Within this framework, six subject-specific parameters were estimated: maximal isometric force of the flexors, maximal isometric force of the extensors, tendon slack length, force–length curve width, parameters governing the nonlinear EMG-to-activation relationship, and a single parameter describing the moment-arm curve, modeled using a cosine-type function. Other model parameters were obtained from a generic OpenSim model (“arm26”; (Seth et al., 2018)) and held fixed. These generic components provided baseline anatomical and physiological relationships, while the estimated parameters captured subject-specific deviations.

For every subject (s), each trial (t), and each muscle (m), the muscle force (F_s,t,m_) was computed as a function of the peak isometric force (*F*_0_*_s_*_,*m*_), the normalized force-length curve (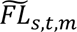), and the activation (*A_s_*_,*t*,*m*_) (Lan, 2002). Note that the force–velocity term is omitted from Equation (1), as the study involved only isometric muscle contractions. In the following equations, an underline notation indicates parameters that were estimated via Bayesian inference.

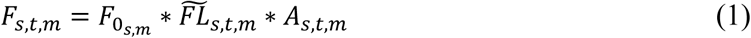

The optimal contractile element (CE) force (*F*_0_*_m_*) was modified for each participant using a Force Scaling (*λ*) parameter that was applied to each of the two muscle groups (flexors and extensors). The extensor and flexor scales apply to all extensor muscles and flexor muscles, respectively.

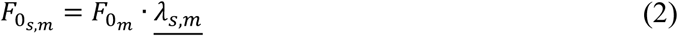

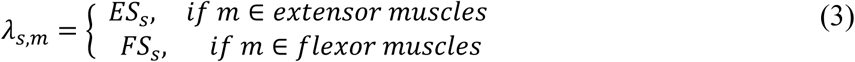

The normalized force-length curve (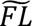) was modeled as an inverted parabola (Lan, 2002) with a fixed width (w) parameter of 0.56 (Figure 2C):

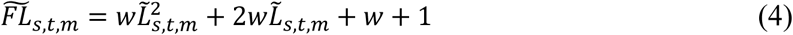

where the normalized muscle length 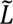 was computed using the length of the muscle tendon (*L_mts_*_,*t*,*m*_), the length of the SEE (*L_Ts_*_,*t*,*m*_) and the optimal muscle length (*L*_0*m*_) as follows:

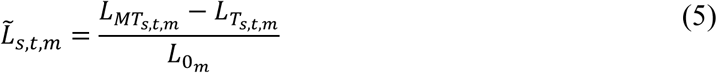

**Figure 2:**
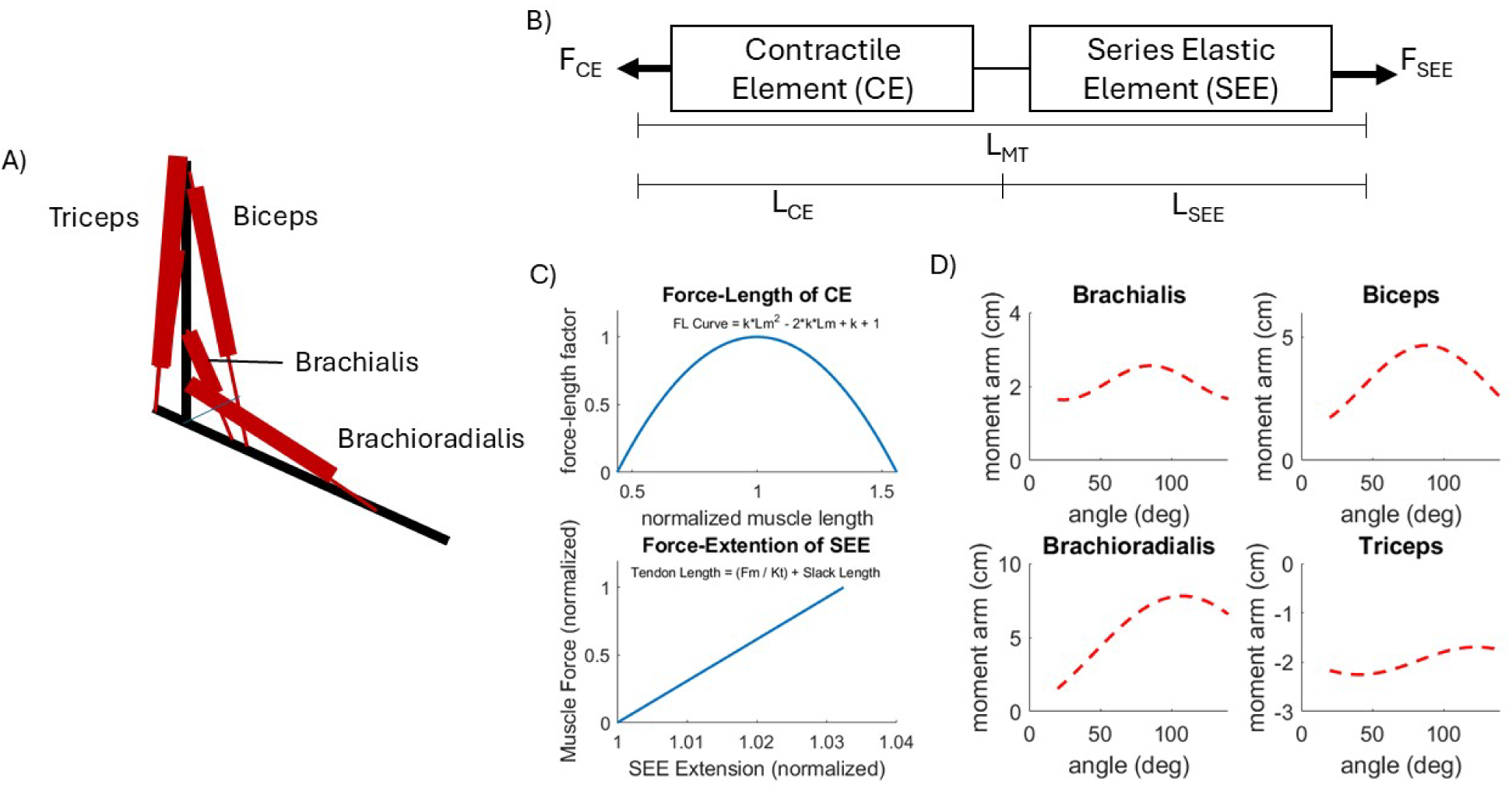
A) The 1-DOF, 5-muscle model: A schematic representation of the musculoskeletal model used for this project. The two triceps muscles have different lengths but share the same moment arm at the elbow, which is why our project shows a single line representing both triceps muscles (Table 1). B) Muscle model: Our simple Hill-type muscle model consists of two elements in series with each other: the series elastic element (SEE) and the contractile element (CE). The length of the SEE plus the length of the CE determined the length of the muscle-tendon unit. C) The force-length relationship of the CE was modeled as an inverted parabola, with a normalized peak isometric force at the optimal CE length. The force-extension of the SEE was modeled as a linear spring, with the SEE stretching by 3.3% of its slack length when the muscle-tendon unit is producing peak isometric force. E) Moment arm curves for each of the muscles in the model for the generic model. These values get scaled based on anthropometrics and altered via the Wave Scale parameter.

**Table 1:** Model parameters from the Generic OpenSim Model.

| **Muscle** | **Peak CE**<br>**Isometric**<br>**Force (N)** | **Optimal CE**<br>**Length (m)** | **SEE Slack**<br>**Length (m)** | **SEE Stiffness**<br>**(N/m)** | **L<sub>MT</sub> at Elbow = 0**<br>**deg (m)** |
| --- | --- | --- | --- | --- | --- |
| Biceps Brachii | 1060 | 0.13 | 0.19 | 1696 | 0.34 |
| Brachioradialis | 990 | 0.11 | 0.13 | 638 | 0.33 |
| Brachialis | 276 | 0.09 | 0.05 | 5693 | 0.15 |
| Triceps Long Head | 800 | 0.13 | 0.14 | 1721 | 0.28 |
| Triceps Lateral<br>Head | 1250 | 0.11 | 0.098 | 3924 | 0.17 |

The total muscle-tendon length (L_MT_) was defined for each condition based on the current elbow angle using a linear function that relates muscle-tendon length to elbow angle. For the L_MT_, the parameters were the length of the muscle-tendon at 0 degrees of elbow flexion (L_MT0_) and the rate of change of the L_MT_ as the elbow flexes (*rθ_t_*) defined based on muscle tendon lengths computed from the arm26 model in OpenSim.

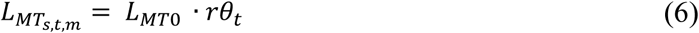

Finally, the mapping of measured EMG to CE activation was represented by a nonlinear power function (Equation 6; (Woods & Bigland-Ritchie, 1983)). We first tested the exponential activation model (Buongiorno et al., 2018; Lloyd & Besier, 2003), but it incurred excessive computational burden and poor Markov chain convergence. Thus, the power function was selected as a mathematically efficient alternative that maintains the physiological curve while facilitating faster and more reliable convergence of the Markov chains.

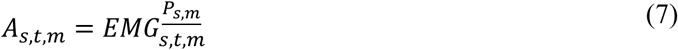

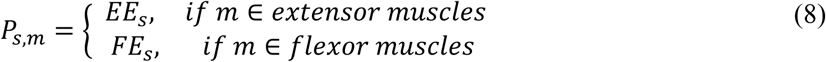

where the muscle activation of the muscle is computed based on the electrical signal measured via filtered and rectified EMG and the non-linear power scale (*P_s_*_,*m*_). The values of *P_s_*_,*m*_ range from 0 to 1, with greater values indicating a more linear relationship between activation and EMG.

The series elastic element (SEE) parameters were slack length (L_slack_) and stiffness (*_kTm_*), which defined the length of the SEE (L_T_) as a function of muscle force (*F_s_*_,*t*,*m*_). SEE stiffness was modeled linearly (Figure 2C) and was assumed to be such that SEE extends by 3.25% of slack length at peak isometric force (Seth et al., 2018). The tendon slack lengths for each muscle were adjusted by scaling the generic tendon slack length using a parameter (*β_s_*).

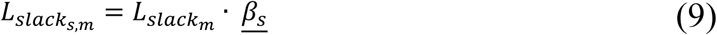

The moment arms (MA) of each muscle were modeled using a cosine-based function derived from measured moment arm data from ten cadaveric specimens ((Murray et al., 2002); Figure 2D). The moment arm magnitudes were also adjusted during the Bayesian inference process by modifying the shape of the cosine wave for each muscle using a parameter (*ψ_s_*).

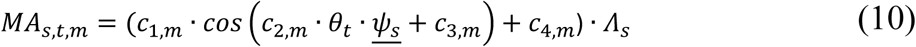

Where c_1_, c_2_, c_3_, and c_4_ are coefficients of the cosine function derived from the measured moment arm data, and *Λ_s_* is a parameter that adjusts the moment arm (y-intercept) based on the participant’s height. The *ψ_s_* is a subject-specific parameter that adjusts the period of the cosine wave. A greater *ψ* value compresses the cosine wave, while a lower *ψ* value stretches the wave. Including the *ψ* parameter allows the model to capture individual differences in joint geometry and muscle attachment points. The scaling factors serve as multipliers to adjust the generic values in Table 1, either increasing or decreasing them.

Finally, the torque produced by the model was computed as:

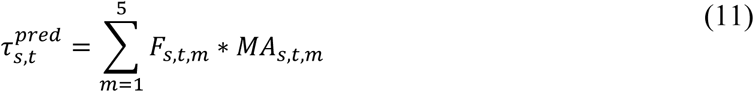

### 2.3 Human Subject Bayesian Parameter Estimation

#### Hierarchical Bayesian Model

The Bayesian model estimated six parameters per subject:

1. *FS_s_*: scaling factor for flexor muscle forces
2. *ES_s_*: scaling factor for extensor muscle forces
3. *β_s_*: scaling factor for tendon slack length
4. *ψ_s_*: scaling factor for the moment arm cosine function (wave scale).
5. *FE_s_*: determines the non-linearity of the EMG to force relationship for the flexors
6. *EE_s_*: defines the non-linearity of the EMG to force relationship for the extensors

To estimate the subject-specific and group-level muscle model parameters from the experimental EMG and torque data, we implemented a hierarchical Bayesian model using Markov Chain Monte Carlo (MCMC) sampling via the Just Another Gibbs Sampler (JAGS) engine in R (Gelman et al., 2013). The human subject dataset was input into the model, along with subject-specific muscle-tendon parameters scaled based on the participant’s standing height (optimal fiber length, tendon stiffness, force-length relationship, angle-dependent LMT, and muscle moment arms).

These parameters were used to compute predicted torque values from EMG inputs via the Hill-type muscle model. The likelihood function was defined based on the measured torque from the dataset 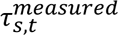 as a Gaussian distribution:

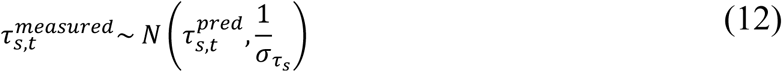

Where 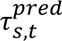 is the predicted elbow torque computed from EMG inputs and model parameters, and 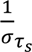 is the precision (inverse of variance) of the torque measurement noise.

Each subject-level parameter was assigned a normal prior with group-level mean and precision. Group-level hyperparameters (means and precision) were assigned to each of the six estimated model parameters (Table 2). Hyperparameters were defined using weakly informative normal and gamma distributions, with truncation bounds to ensure physiological plausibility.

**Table 2.** The parameters, priors, and hyper-priors for each parameter in the Bayesian model. *N*_(*x*,*y*)_indicates a lower bound of x and an upper bound of y.

| Parameters | Priors | Hyper-Priors |
| --- | --- | --- |
| Extensor Force Scale | $ES_s \sim N_{(0.1,+\infty)}(\mu_{ES}, \frac{1}{\sigma_{ES}})$ | $\mu_{ES} \sim N_{(0.01,+\infty)}(1,0.1)$<br>$\sigma_{ES} \sim InvGamma(0.1,0.1)$ |
| Flexor Force Scale | $FS_s \sim N_{(0.1,+\infty)}(\mu_{FS}, \frac{1}{\sigma_{FS}})$ | $\mu_{FS} \sim N_{(0.01,+\infty)}(1,0.1)$<br>$\sigma_{FS} \sim InvGamma(0.1,0.1)$ |
| Tendon Slack Length Scale | $TSL_s \sim N_{(0.8,1.7)}(\mu_{TSL}, \frac{1}{\sigma_{TSL}})$ | $\mu_{TSL} \sim N_{(0.8,1.3)}(1,0.01)$<br>$\sigma_{TSL} \sim InvGamma(80,0.1)$ |
| Moment Arm WaveScale | $\psi_s \sim N_{(0.5,1.5)}(\mu_\psi, \frac{1}{\sigma_\psi})$ | $\mu_\psi \sim N_{(0.01,1.5)}(1,0.01)$<br>$\sigma_\psi \sim InvGamma(80,0.1)$ |
| Extensor Nonlinear Exponent | $EE_s \sim N_{(0.5,1)}(\mu_{EE}, \frac{1}{\sigma_{EE}})$ | $\mu_{EE} \sim N_{(0.1,+\infty)}(0.7,1)$<br>$\sigma_{EE} \sim InvGamma(100,0.1)$ |
| Flexor Nonlinear Exponent | $FE_s \sim N_{(0.5,1)}(\mu_{FE}, \frac{1}{\sigma_{FE}})$ | $\mu_{FE} \sim N_{(0.1,+\infty)}(0.7,1)$<br>$\sigma_{FE} \sim InvGamma(100,0.1)$ |
| Data | Likelihood | Hyper-Priors |
| $\tau_{measured}$ | $N\left(\tau_{s,t}, \frac{1}{\sigma_\tau}\right)$ | $\sigma_\tau \sim InvGamma(0.1,0.1)$ |

We applied the hierarchical Bayesian inference framework to estimate subject-specific muscle model parameters from the human EMG and torque data. The model’s performance was evaluated using the root-mean-squared error, mean absolute error, and mean absolute percentage error between the predicted and measured elbow torque values. We assessed convergence using the updated Gelman-Rubin R-hat statistics (Vehtari et al., 2021) and visual inspection of trace plots, which were used to confirm stable sampling and good mixing across chains. Posterior means and 95% credible intervals were computed for each parameter for each subject.

#### Comparison of Models with Different Parameter Combinations

Testing different model alternatives allows us to test whether the parameters in the 6-parameter model improve predictive performance, or whether a more parsimonious model performs equally well, or better. More complex models are only warranted when they deliver demonstrably better predictive performance. Selecting the model with the lowest WAIC ensured that our main analysis was based on a parameterization that balanced goodness of fit with generalizability. We tested different combinations of model parameters to assess which set best fits the experimental data. To achieve this, we tested three other models with different numbers of free parameters and evaluated them based on the Watanabe-Akaike information criterion (WAIC) values (Watanabe, 2013). Besides the six-parameter model, we also tested a 3-parameter model (Flexor Scale, Extensor Scale, Tendon Slack Scale), a 4-parameter model (Flexor Scale, Extensor Scale, Tendon Slack Scale, Wave Scale), and a 7-parameter model (Everything in the 6-parameter model, except the Tendon Slack Scales split into two parameters, one for the flexor muscles and one for the extensor muscles).

All models were fit using jags.parallel() with the following settings: Iterations=16000, Burn-in=8000, Thinning=10, Chains=4, Seed=42. Posterior samples were extracted for all individual and group-level parameters, including the predicted torque values.

#### Comparison of Different Parameter Estimation Methods Through Leave One Subject Out Fewshot Learning

We next compared the strength of the hierarchical Bayesian model approach compared to (i) a point-estimate method using MATLAB’s constrained nonlinear optimization function *fmincon* (MATLAB R2025a, The MathWorks Inc., Natick, MA, USA) with a maximum-likelihood objective, and (ii) a non-hierarchical Bayesian model. For the non-Bayesian model, the parameters were optimized individually using *fmincon*. For the two Bayesian models, predictions were generated using the posterior median of each parameter.

We initially partitioned the data using a Leave-One-Subject-Out (LOSO) scheme, in which data from all subjects except the test subject were included in the training set. To further evaluate model stability and generalization, we then incrementally added increasing fractions of the test subject’s trials to the training set and evaluated torque prediction accuracy on the remaining held-out trials. This approach provides a systematic way to evaluate how model performance changes as subject-specific data are incorporated and is consistent with established practices for assessing hierarchical Bayesian models (Ahuja & Sethia, 2024). To quantify how accurately each modeling approach predicted torque on held-out trials, we used mean percentage error (MPE). MPE expresses the prediction error as a percentage of the true measured torque, allowing performance to be compared across subjects and conditions with different absolute torque magnitudes. Because MPE normalizes errors relative to the observed torque magnitude, it provides an interpretable, scale-free metric suitable for comparing the hierarchical Bayesian model, the non-hierarchical Bayesian model, and the non-Bayesian *fmincon*-based fit across progressively larger training-set sizes.

#### Validation with the Other Arm

To evaluate the robustness of our experimental results, we also used the best-fitting model to perform Bayesian inference for the seven subjects who underwent bilateral testing. In this study, young, healthy adults are expected to exhibit symmetrical musculoskeletal and tendon properties between limbs, such that within-subject differences between the left and right sides should be relatively small compared to between-subject differences. Therefore, we incorporated arm side as a covariate in the hierarchical priors. As an example, the subject-level Flexor Force Scale was modeled as:

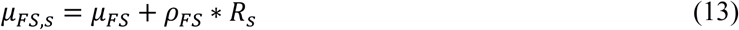

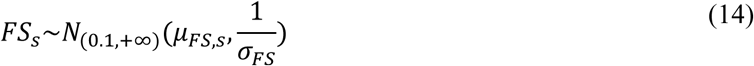

Where *ρ_FS_* is a group-level regression coefficient for arm side, and *R_s_* is a subject-level indicator covariate dictating the dominant or non-dominant arm (1 if the arm is on the dominant side).

#### Sensitivity Test on Prior Distributions

To test the robustness of our results to a range of prior distributions, we conducted a prior sensitivity analysis (Supplement 1) by rescaling the priors. For normally distributed priors, the standard deviation was multiplied by factors ranging from 0.1 to 10 (10 log-spaced values). Similarly, for gamma-distributed priors, both the shape and scale parameters were multiplied by the same set of log-spaced factors. Varying the priors allows us to verify that the model’s conclusions are driven by the data rather than by arbitrary prior choices. By rescaling prior widths over several orders of magnitude, we test whether posterior estimates for both group-level and individual-level parameters remain stable across a broad range of plausible assumptions. The hierarchical Bayesian model was refitted under each specification; posterior medians for group-level and individual-level parameters are reported in Supplementary Tables S1.1–S1.2 as percentage changes relative to those reported in the main manuscript.

#### Sensitivity Test on Co-activation

We noticed that some participants exhibited some levels of co-contraction. Thus, to assess the sensitivity of our parameter estimates to EMG co-activation, we conducted a separate analysis in which we removed the co-activation EMG from the antagonist muscles and re-fit the hierarchical Bayesian model using these modified inputs. Removing antagonist co-activation EMG allowed us to evaluate whether our parameter estimates were influenced by residual or misattributed muscle activity. Because co-activation can artificially alter inferred torque contributions, repeating the analysis with these signals removed helped confirm that the model’s parameter estimates reflect true muscle–tendon dynamics rather than artifacts of EMG preprocessing. The results of this sensitivity analysis are presented in Supplementary 2.

### 2.4 Synthetic Data

#### Synthetic Data Generation

We validated our Bayesian estimation by evaluating our ability to recover known parameter values in a synthetic dataset with characteristics similar to those from our experimental data. Using MATLAB, we generated isometric elbow torque and corresponding EMG signals across a sample of 50 synthetic subjects to evaluate the performance of the Bayesian inference algorithm in estimating muscle model parameters. For each of the 50 subjects, we used the posterior median parameter estimates derived from the experimental data to sample subject-specific values for maximal isometric force (CE property), force–length curve width (CE property), tendon slack length (SEE property), the parameters governing the nonlinear EMG-to-activation relationship, moment arms, and muscle–tendon lengths across three elbow angles (20°, 70°, and 120°). To account for individual anatomical differences, musculoskeletal model parameters were scaled using anthropometric data, generated from the distribution of participant heights in our subject pool. Moment arm profiles were computed using a cosine-based function, with subject-specific scaling derived from measured height relative to our reference model. The individual parameters for the simulated subjects were sampled as: *FS_s_*∼*N*(2.1, 0.6^2^), *ES_s_*∼*N*(1.9, 0.6^2^) *TSL_s_*∼*N*(1.1, 0.06^2^), *ψ* ∼*N*(0.86, 0.15^2^), *P_F_*_,*s*,*m*_∼*N*(0.66, 0.06^2^), *P_E_*_,*s*,*m*_∼*N*(0.77, 0.06^2^), *MAScale_s_*∼*N*(1.1, 0.1^2^).

Each artificial subject was evaluated using the same protocol as for the experimental participants. There were six conditions in total: three angles and two directions (flexion and extension). For each of the six conditions, the artificial subject generated a peak isometric torque by setting the activation of the agonist muscles to its maximum value (i.e., full activation, A=1), while the activation of the antagonist muscles was set to 0. This allowed us to measure the peak torque capacity at the angle based on the subject-specific muscle-tendon lengths and moment arms. We then performed three submaximal blocks, where we computed the artificial EMG associated with producing torques of 10%, 25%, and 40% of the peak isometric torque for that condition. For each condition and torque magnitude, 10 trials were generated, resulting in a total of 180 trials per artificial subject.

For each trial, target torque values were perturbed by ±10% to simulate the allowable experimental error. Muscle forces were then distributed across the muscles using a constrained optimization (via fmincon) that minimized the sum of each muscle’s EMG raised to the power 3/p to represent the nonlinearity of muscle EMG in the experimental data.

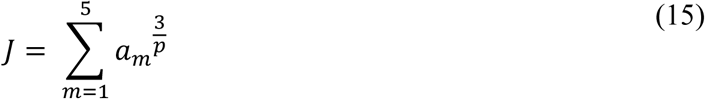

To emulate realistic measurement conditions, Gaussian noise, signal-dependent (SD = 10% of the signal magnitude), was added to the simulated EMG signals. This step is important because real EMG data are inherently noisy due to physiological variability, electrode placement, and signal-dependent noise (Clancy et al., 2002; De Luca et al., 2010; Huigen et al., 2002). Incorporating noise ensures that the synthetic data reflect experimental uncertainty, prevents overfitting to idealized signals, and tests whether the Bayesian model can robustly estimate parameters under realistic conditions. The 10% noise level was chosen to match the noise from our experimental dataset.

The simulation produced EMG signals for all five muscles and measured torque for each of the 50 artificial subjects, based on each subject’s unique combination of muscle parameters and moment arm profiles.

#### Bayesian Parameter Recovery

We applied the hierarchical Bayesian inference framework described in Section 2.3 to estimate subject-specific parameters of the muscle model from synthetic EMG and torque data. For each of the 50 synthetic participants, we used EMG signals and torque measurements to recover the same six parameters that are evaluated from actual data (Flexor Scale, Extensor Scale, Tendon Slack Scale, Wave Scale, Extensor Non-Linear Power, and Flexor Non-Linear Power). We also computed correlation coefficients between pairs of parameters to assess their correlation across iterations of the MCMC algorithm (*Supplement 3*).

Additionally, to investigate the influence of antagonist co-activation on parameter recovery, we generated a second synthetic dataset with 50 synthetic participants in which antagonist muscle activations increased with torque level. To model this relationship, linear regressions between antagonist EMG amplitude and torque were performed for each experimental participant. The resulting regression coefficients (slope and intercept) were used to estimate population distributions, from which subject-specific parameters were sampled during simulation. Antagonist EMG activations were then generated according to the sampled torque-dependent relationships. Co-activation was implemented by enforcing a nonzero lower bound on antagonist muscle activation during the constrained optimization procedure (fmincon), thereby ensuring simultaneous activation of agonist and antagonist muscles.

## 3. Results

### 3.1 Experimental Results: Parameter Estimation and Model Performance

#### Surface EMG-torque Relationship

Participants performed isometric elbow flexion and extension across a range of joint angles and target torque levels. A representative example of the relationship between EMG amplitude and elbow joint torque for the five recorded muscles from a single subject is shown in Figure 3A. Overall, agonist EMG amplitude exhibited an approximately linear relationship with net elbow torque, as previously described (Shin et al., 2009). However, some individuals co-activated the flexors and extensors, as shown by asymmetrical “V-shape” EMG-Torque curves (Supplement 4).

**Figure 3:**
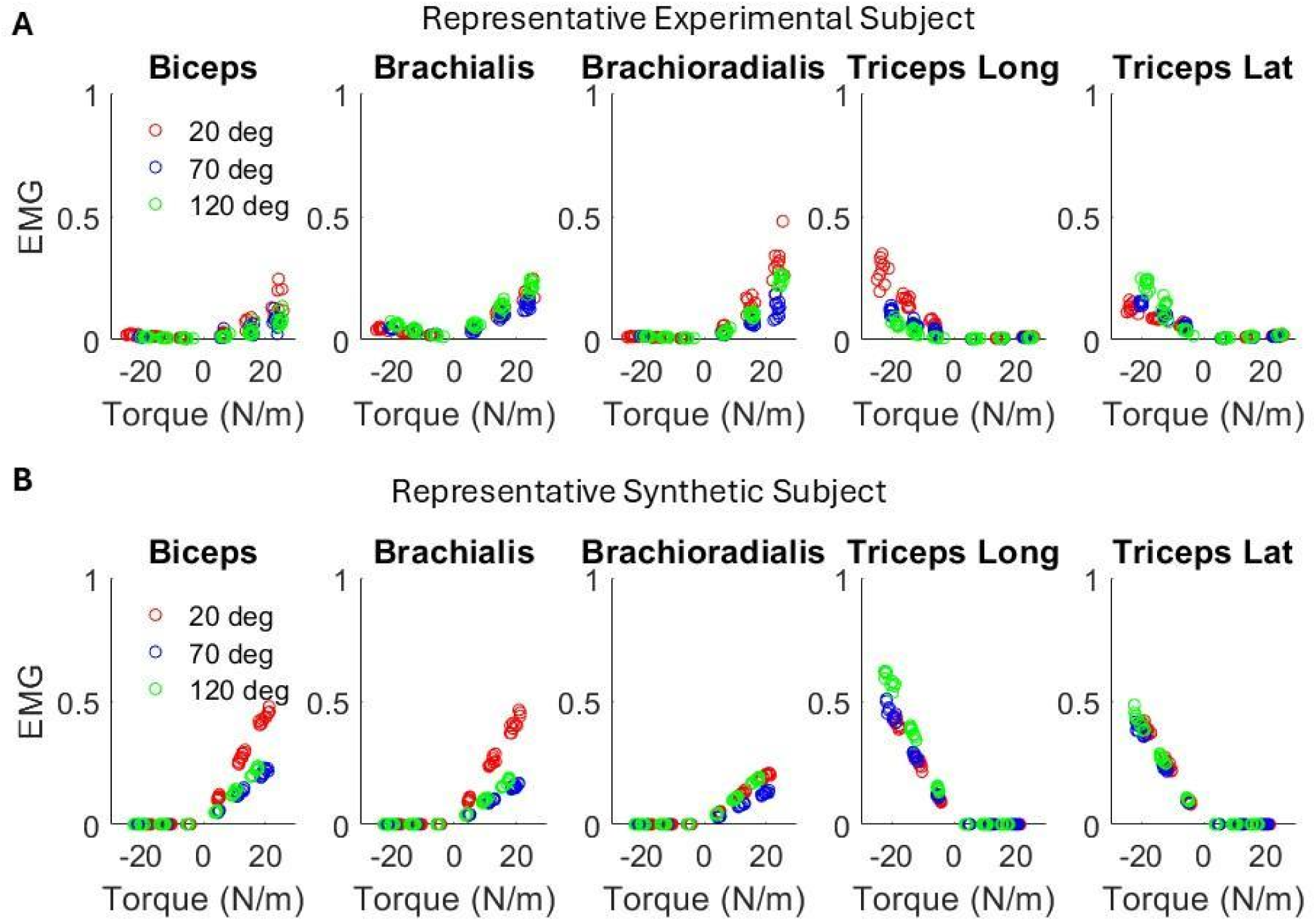
Surface EMG-Torque. A) The surface EMG versus elbow joint torque for a representative experimental subject. B) Simulated EMG and torque relationship for a synthetic participant. The y-axis is the average amplitude of the normalized surface EMG (normalized by MVC) within each trial, and the x-axis is the measured net elbow joint torque (Extension torques are plotted as negative values) for each trial.

**Table 3:**
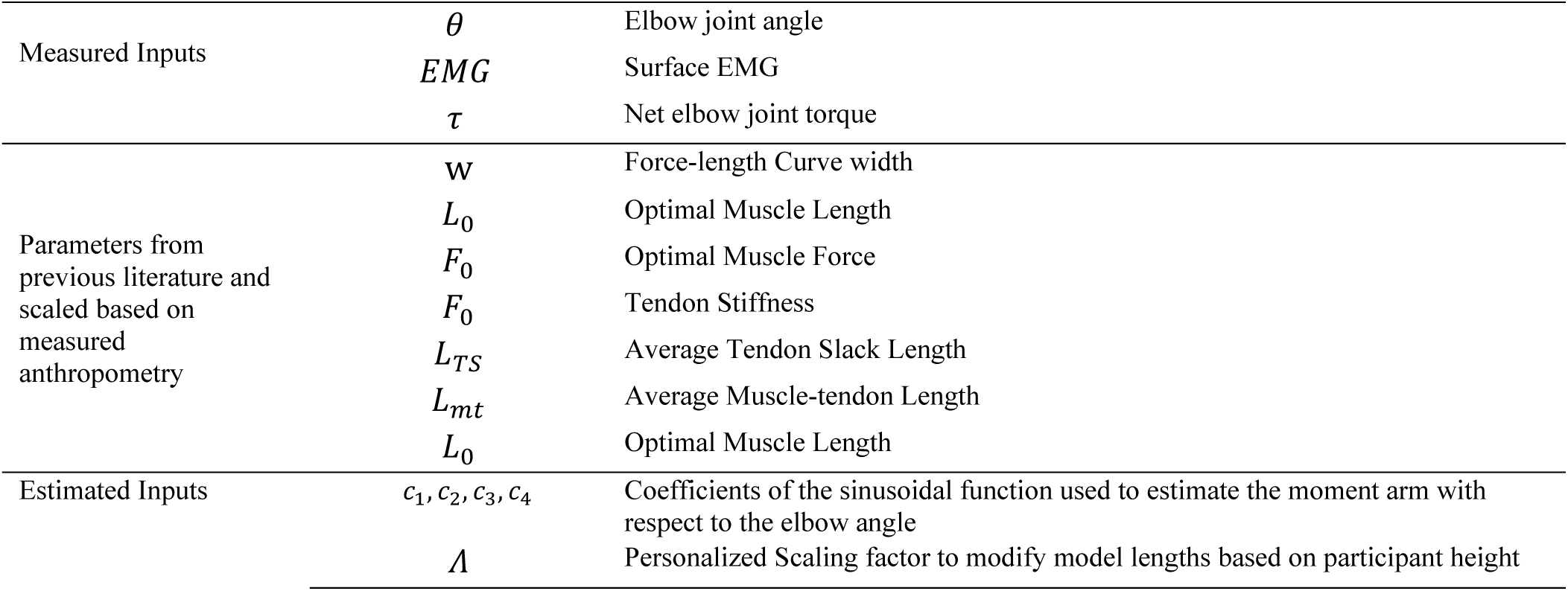
Model parameter names and symbolic representations: Our data collection process recorded elbow joint angles, surface electromyography (EMG), and net elbow joint torque (Measured Inputs). The unscaled model values were recorded from prior literature estimates (Holzbaur et al., 2005; Murray et al., 2002; Seth et al., 2018)

#### Hierarchical Bayesian estimation of Hill Model fits and comparison

To evaluate the suitability of our hierarchical Bayesian framework for empirical data, we fitted the model to EMG and torque measurements collected from 14 healthy adults performing isometric elbow flexion and extension tasks.

We compared different model configurations using torque fitting error, representing predictive accuracy, and WAIC, a measure of model fit (*Supplement* 6). Compared to the 3 and 4-parameter models, the 6-parameter model performed best, yielding the highest coefficient of determination (R² = 0.96) and the lowest WAIC value (Table 4). The 7-parameter version, in which the Tendon Slack Scale was estimated separately for flexors and extensors, was excluded from the comparison because it produced physiologically implausible, negative muscle forces. Negative muscle forces arise when the CE lengths fall beyond the valid range of the inverted-parabolic force-length curve, where the curve is not truncated at zero, thus it can yield negative muscle forces.

**Table 4:**
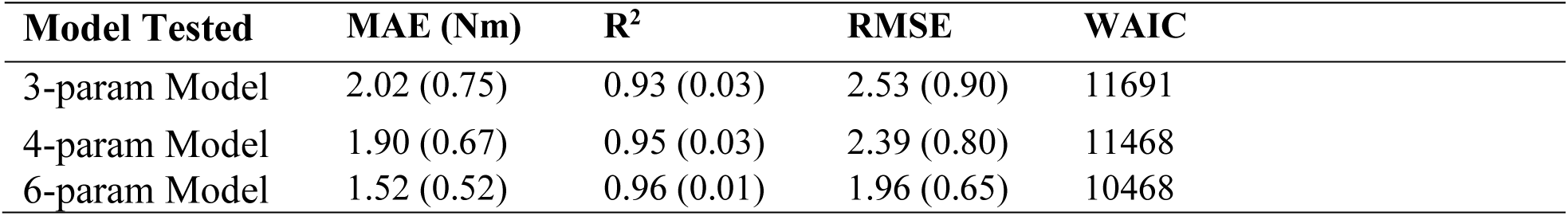
Model fits for each of the three types of models we tested. The “3-param Model” includes the Flexor Scale, Extensor Scale, and Tendon Slack Scale. The “4-param Model” includes the 3-param model plus the Wave Scale. The “6-param Model” includes everything in the 4-param model and the Extensor Power Scale and Flexor Power Scale. MAE = mean absolute error, RMSE = root mean squared error, WAIC= Watanabe-Akaike Information Criteria.

Across all fitting trials and for each subject, predicted versus measured torque in the 6-paramter model showed high agreement, with MAE values below 5% of peak torque (Supplementary S5.1). Error histograms revealed no systematic bias, and cross-validation on held-out trials (see Methods) confirmed robust generalization (Figure 4). All 4 chains achieved satisfactory convergence, with R-hat values below 1.02 for all parameters and trace plots showing good mixing. Total computational time for full hierarchical sampling was approximately 2 hours on an 8-core machine.

**Figure 4:**
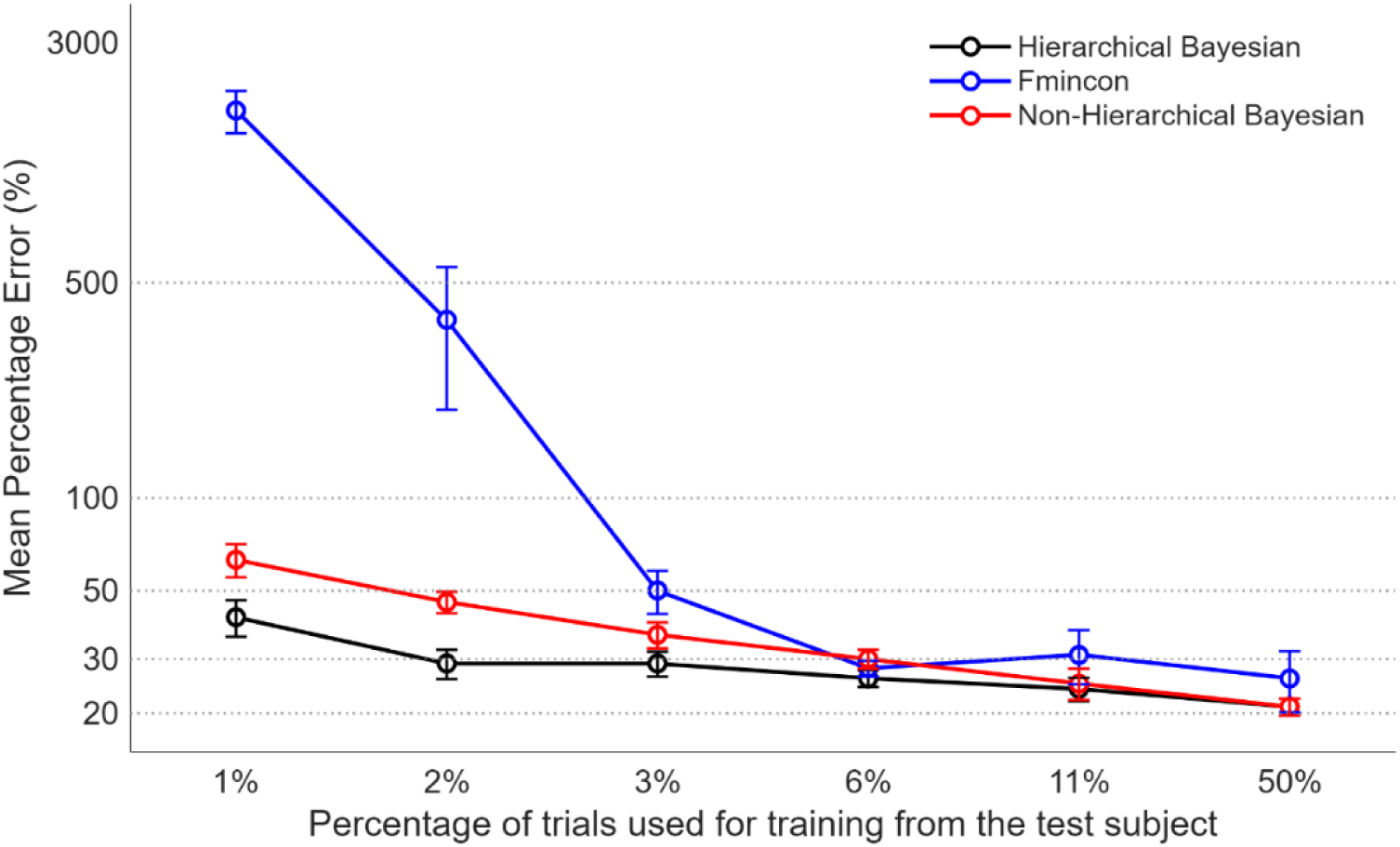
The mean percentage force reconstruction error of the leave one out cross validation procedure using different parameter estimation methods, with varying amounts of data that were left out for a given participant.

#### Posterior Inference of Model Parameters

The posterior distributions for all parameters (Figure 5) indicate both subject-level variability and population-level central tendencies. Subject-specific estimates display substantial uncertainty, reflecting differences across individuals while demonstrating the hierarchical model’s capacity to provide regularized parameter estimates. At the population level, the posterior means and their associated intervals summarize overall tendencies across the cohort.

**Figure 5:**
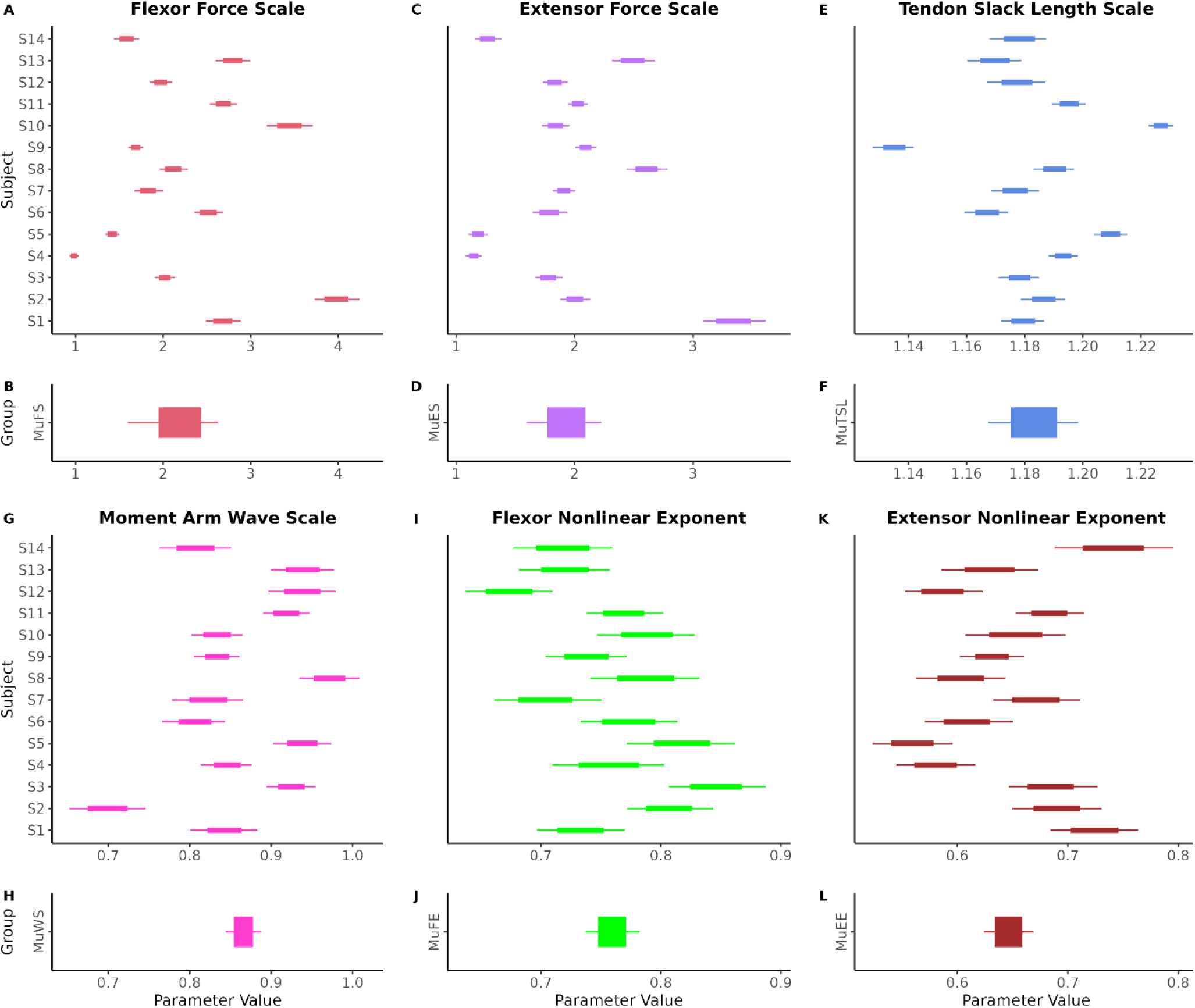
Individual and group level posterior estimates for six model parameters: Flexor Force Scale, Extensor Force Scale, Moment Arm Wave Scale, Tendon Slack Length Scale, Flexor Nonlinear Exponent, and Extensor Nonlinear Exponent. In the top panel, horizontal bars represent the posterior mean and 95% highest density interval (HDI) for each subject, highlighting inter-individual variability and estimation uncertainty. The bottom panel displays the posterior group means and associated uncertainty intervals, representing the central tendency and variability across the population.

#### Muscle Force Scale

The group-level posterior means for the flexor and extensor force scaling parameters were approximately 2.0, suggesting that participants were, on average, twice as strong as the generic OpenSim “arm26” musculoskeletal model. The individual-level posterior estimates for these force-scaling parameters ranged from approximately 1.0 to 3.5 across participants, indicating substantial variability in muscle strength across participants and demonstrating the model’s ability to distinguish subject-specific characteristics.

#### Tendon Slack Scale

In contrast, the recovered tendon slack length scaling parameters exhibited a relatively narrow range (approximately 1.14–1.22), indicating limited inter-subject variability. To assess the validity of the tendon slack length estimates, normalized muscle lengths were computed using Equation (5), in which the Tendon Slack Scale was the only estimated parameter, and all other parameters were derived from the generic OpenSim “arm26” model. These normalized muscle lengths were then mapped onto the muscle force–length relationship to obtain the corresponding force–length (F-L) factors, and the resulting distributions for each subject were overlaid onto the theoretical F-L curve (Figure 6A-E). The model produced physiologically plausible force–length factors, with the contractile element operating primarily on the ascending limb. Further validation was performed by comparing the normalized force–length curves with previously reported cadaveric data (Murray et al., 2000) (Figure 6F-J). The range of muscle lengths observed across the three tested postures generally overlapped with that observed in the cadaver study (Murray et al., 2000), supporting the validity of our estimated Tendon Slack Length. For example, the estimated length of the biceps ranged from 0.5 to 1.0, consistent with empirically measured values. However, the normalized muscle length for the triceps longus was slightly smaller in our study. This discrepancy may be attributed to measurement noise and/or the smaller minimum elbow angle used in the current protocol (20° vs. 30°).

**Figure 6:**
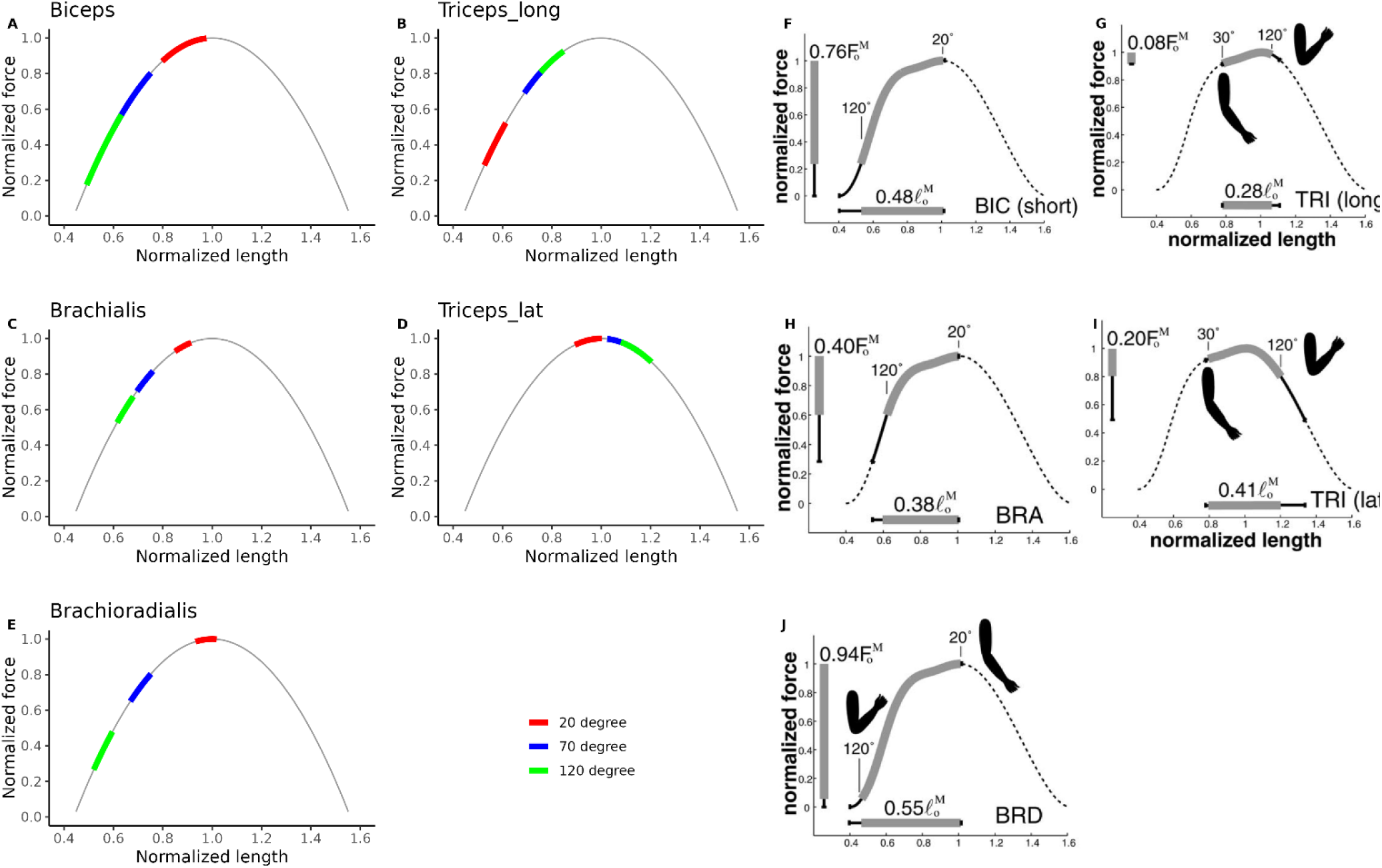
Force-Length Curves: The colored bars show where on the force-length curve the contractile element operates for each of the angle conditions in the 6-parameter model. Four of the five muscles investigated primarily operate on the ascending limb of the force-length curve, while the triceps lateralis seems to operate at the apex of the force-length curve. For comparison, force–length curves from a previous cadaver study (Murray et al., 2000) are shown on the right.

#### Muscle Moment Arm Characteristics

The moment arm scaling factors ranged from approximately 0.7 to 1.0, indicating that the cosine-shaped moment arm curves were stretched (i.e., widened). The range reflects subject-specific differences in moment arm geometry captured by the scaling parameter (Figure 5G).

#### Non-linear EMG to Activation Relationship

Finally, the hierarchical posterior estimates for the nonlinear EMG–activation exponents were consistently less than one for both elbow flexors and extensors (median ≈ 0.76 for flexors and ≈ 0.65 for extensors; Figure 5I–L). This sublinear behavior reflects the concavity of the power function in Equation (6), whereby increases in EMG amplitude yield diminishing returns to activation at higher input levels. Consequently, torque production scales nonlinearly with EMG amplitude, increasing less proportionally in the higher activation range

#### Posterior Parameters from both arm sides

Posterior distributions of individual-level parameters were estimated separately for the left and right sides in a subset of seven participants (Figure 7). Posterior means and 95% credible intervals were computed for all parameters. To evaluate inter-limb symmetry and quantify the effect of hand dominance, posterior estimates were compared between sides, with arm laterality included as a group-level covariate. Overall, the distributions for the left and right sides showed substantial overlap. Within-subject variability (left vs. right) was notably smaller than between-subject variability, consistent with expectations. The 95% credible intervals for the group-level coefficients included zero, indicating largely symmetrical parameter estimates. However, more than 87% of the posterior coefficients for the Extensor Force Scale were greater than zero, suggesting a possible bias towards stronger extensors in the dominant arm.

**Figure 7:**
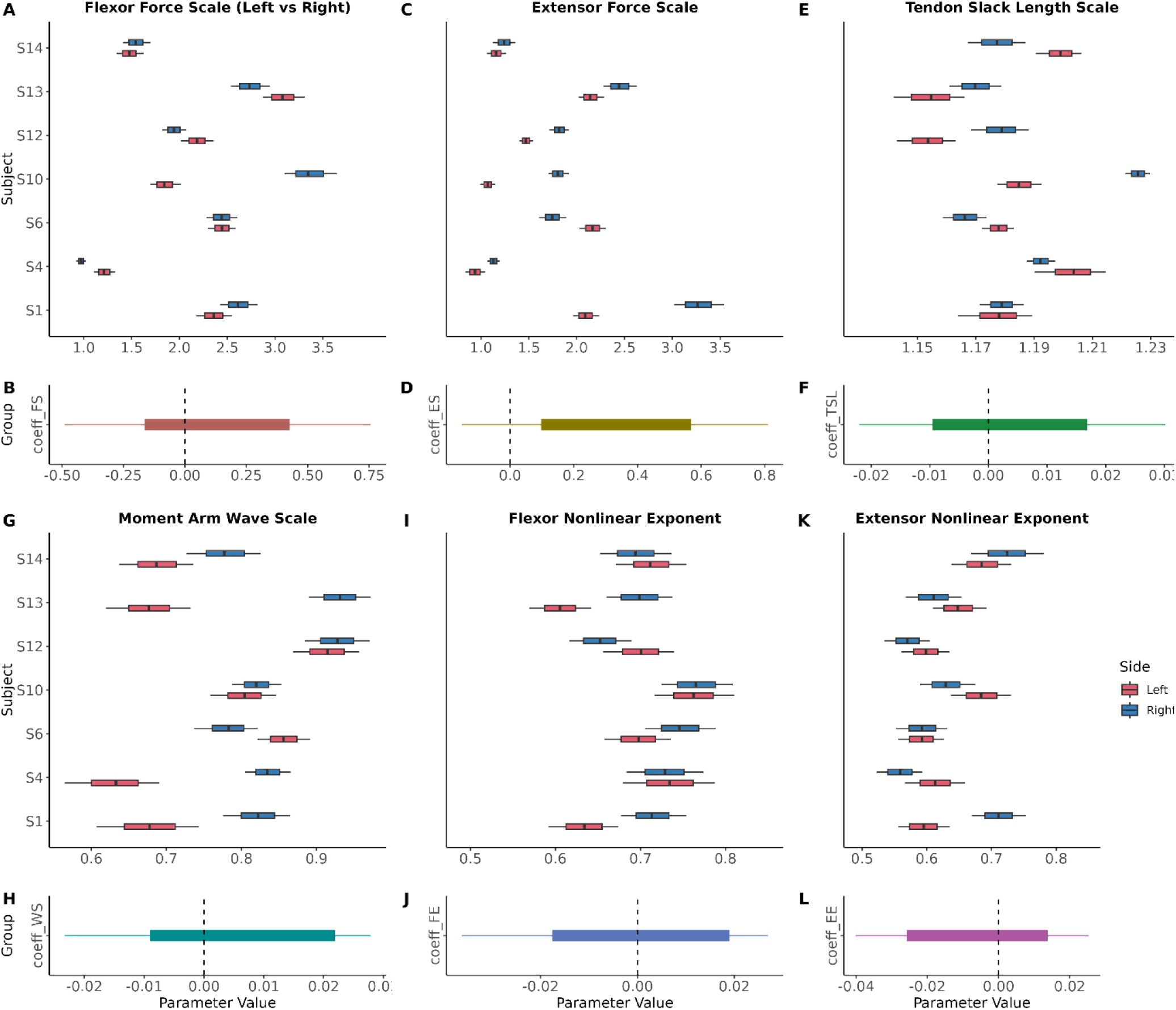
Posterior distributions of the six left and right-side individual parameters, and the posterior distributions of the covariate coefficients for the six model parameters.

#### Sensitivity Test on Co-activation

As shown in Supplementary S2.1 and S2.2, removing co-activation from the antagonist muscles led to substantial changes in posterior estimates of parameters such as Flexor Scale, Extensor Scale, Flexor EMG Exponent, and Extensor EMG Exponent, whereas estimates of Tendon Slack Length Scale and Wave Scale were comparatively less affected. At the individual level, subjects with low co-activation indices (e.g., Subjects 1, 12, and 14) exhibited smaller changes in the Flexor and Extensor Scales following removal of antagonist EMG compared to subjects with high co-activation indices (e.g., Subjects 4, 5, and 6). These findings suggest that neglecting EMG co-activation can significantly alter the inferred EMG–torque relationship, consistent with previous reports (Brown & McGill, 2008).

#### Sensitivity to the prior distribution

To assess the robustness of parameter estimation to prior specification, we performed a prior sensitivity test by rescaling the priors (Supplement 1). As shown in Table S1.1, percentage changes in all group-level means remained within 5% of the baseline, indicating that the posterior group-level parameter medians are insensitive to prior choice. For the group-level standard deviations, the percentage changes for the Force Scales and Tendon Slack Scale are below 20%. Although the percentage changes for Wave Scale and the two Exponents exceed 50%, their absolute differences across different prior conditions are small (all below 0.05), suggesting minimal practical impact.

Table S1.2 summarizes the percentage change in posterior medians of each individual-level parameter across participants. Most changes were within 5%, with only three individual parameters exceeding a 10% difference from the baseline, suggesting that the individual-level parameters are also robust to different prior specifications.

#### Comparison of Hierarchical Bayesian estimation to non-hierarchical and point-estimate methods

We finally evaluated the torque prediction performance with our hierarchical Bayesian model compared to (i) the interior point algorithm in the MATLAB *fmincon* function and (ii) a non-hierarchical Bayesian model (see Methods). Although the results show that mean percentage errors were similar across the three methods when more than 6% of the validation subjects’ trials were used in the training set, the non-Bayesian method exhibited much larger errors and greater variability (standard error) when fewer than 6% of the trials were included (Figure 4). This pattern suggests that Bayesian methods are more stable and generalize better to unseen data, or when data from the test subject is limited. Additionally, when comparing the two Bayesian models, the hierarchical model generally performed better (lower error and standard error) when less than 11% of the trials were used for training, indicating that group-level parameters help improve predictions under sparse-data conditions.

### 3.2 Synthetic Results: Parameter Recovery and Model Validation

To evaluate the effectiveness of our Bayesian framework in recovering true muscle model parameters, we generated a synthetic dataset for 50 synthetic participants (Figure 3B).

#### Model Fitting and Convergence

Model fitting to the synthetic data showed excellent performance, with an R² of 0.99, an RMSE of 1.26, and an MAE of 0.89. All Markov chains met convergence criteria, with R-hat values below 1.05, well within the acceptable threshold of 1.10 (Gelman et al., 2013; Vehtari et al., 2021), indicating reliable posterior estimation.

#### Parameter Recovery with the Synthetic Data

To assess parameter recovery accuracy, posterior means were compared with the ground-truth parameter values across all synthetic subjects. Scatter plots revealed tight clustering along the line of identity for all parameters (Figure 8), with average percent errors consistently falling below 5%. The Extensor Force Scale (mean error = 3.0%, max = 8.3%) and Flexor Force Scale (mean error = 4.1%, max = 10.8%) exhibited similar error magnitudes, while the Tendon Slack Length Scale demonstrated the highest accuracy (mean error = 1.0%, max = 4.2%). Additionally, the hierarchical posterior distributions at the population level were centered near the true parameter values (Figure 9), supporting the model’s accuracy.

**Figure 8:**
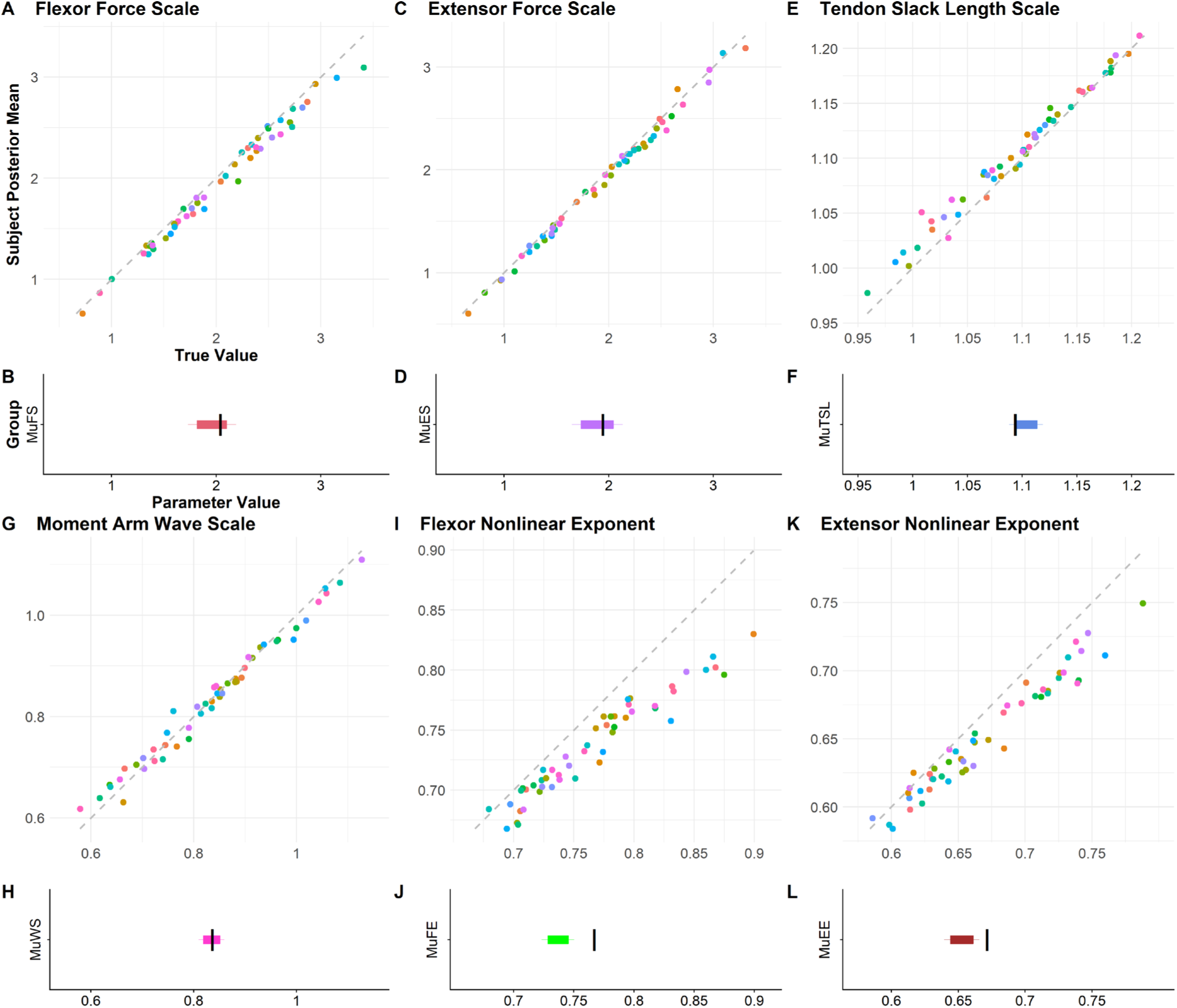
Parameter Recovery for the 6-parameter Model using the synthetic dataset. Top panel shows the individual parameter recovery result. The x-axis is the true value for each subject, set through a sample of a normal distribution for each parameter, and the y-axis is the mean of the posterior distribution of the MCMC result. Each dot is a synthetic subject, and dots along the identity line show that the recovered parameter from the MCMC algorithm is equal to the true parameter. Extensor Scale means percent error = 3.0%, max error = 8.3%; Flexor Scale means percent error = 4.1%, max error = 10.8%. Bottom panel shows the hierarchical parameter recovery results. Each boxplot represents the posterior distribution of the group-level ean for a given parameter, showing the central tendency and variability across subjects. These distributions reflect the model’s ability to capture inter-individual differences while leveraging hierarchical pooling to estimate them. The vertical black bars show the true population mean for each of the six parameters.

**Figure 9:**
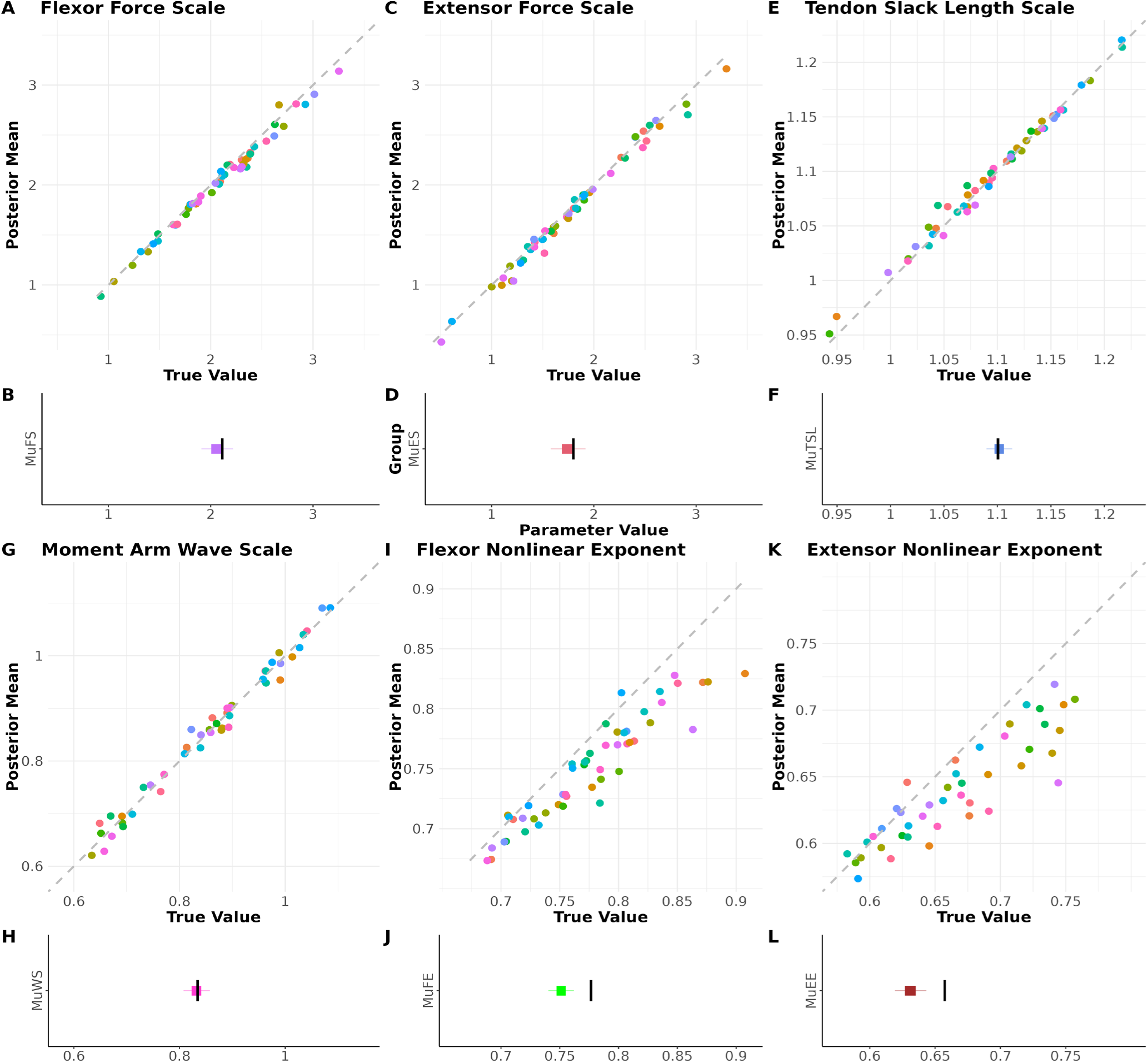
Parameter Recovery for the 6-parameter Model using the synthetic dataset with co-activation from the antagonist muscles.

Similar scatter plots generated using the synthetic dataset with antagonist co-activation are presented in Figure 9. Parameter recovery performance remained comparable to that obtained from the synthetic dataset without co-activation. The Extensor Force Scale (mean error = 3.7%, maximum error = 15.5%) and Flexor Force Scale (mean error = 2.7.%, maximum error = 7.3%) exhibited the largest estimation errors, whereas the Tendon Slack Length Scale was recovered with the highest accuracy (mean error = 0.5%, maximum error = 2.5%). Overall, these results suggest that the proposed parameter estimation framework is robust to the presence of antagonist co-activation, maintaining accurate parameter recovery even when co-activation is incorporated into the synthetic EMG data.

#### Correlation across Posterior Parameters

To examine potential dependencies among the six model parameters, we performed a cross-correlation analysis (*Supplement* 3). The resulting correlation matrix reveals low pairwise correlations between parameters, with no values exceeding an absolute value of 0.31, indicating minimal parameter interdependence within the model.

## 4. Discussion

This study demonstrates the feasibility of using hierarchical Bayesian inference to estimate subject-specific muscle model parameters from EMG and torque data during isometric elbow contractions. By modeling both individual and group-level variability, our approach enables robust recovery of physiologically meaningful parameters, including peak isometric force scaling, tendon slack length, and EMG-to-activation nonlinearity, without relying on imaging or invasive measurements. The framework was applied to experimental data from young, healthy adults and then validated on synthetic data with known ground-truth parameters. Previous work has demonstrated the utility of Bayesian inference for estimating plausible muscle forces from kinematic data, particularly in addressing the muscle redundancy problem and quantifying uncertainty in excitation trajectories (Johnson et al., 2022). We extend this framework to infer the underlying parameters of a muscle model from EMG and torque data, offering a more direct route to physiological personalization.

When applied to experimental data, the model yielded plausible parameter estimates across subjects, with credible intervals that reflected inter-individual variability. The magnitudes of the estimated force scaling factors suggest that our method captured the physiological differences between the young, healthy adults we examined and the cadaveric specimens from which generic model parameters are commonly taken (Buchanan, 1995; Buchanan et al., 2004). The group-level posterior means for the flexor and extensor force scaling parameters were approximately 2.0, indicating that participants were, on average, twice as strong as the generic musculoskeletal model. This finding aligns with expectations, given that the “arm26” model parameters were derived from cadaveric data, often from older individuals with reduced muscle strength. This finding also highlights the importance of individualized parameter estimation and the limitations of relying solely on generic or literature-based values for muscle strength in biomechanical simulations (Rajagopal et al., 2016). Notably, the individual-level posterior estimates for these force-scaling parameters ranged from approximately 1.0 to 3.5 across participants, suggesting that the model discriminated among individuals with variation in muscle strength. In contrast, the recovered tendon slack-length scaling parameters showed a much narrower range, from approximately 1.14 to 1.22, indicating limited variation between participants. This may reflect either a true physiological consistency in tendon slack length among healthy young adults or a limitation in the model’s sensitivity to detect variation in this parameter from surface EMG and torque data alone. The estimated moment arm scaling factor provides a simple but informative means of capturing subject-specific deviations from generic geometry. The fact that values were generally below unity suggests that the cosine-based template derived from Murray’s data (Murray et al., 2002) tends to overestimate the angular breadth of moment arm variation in our cohort, requiring a stretching of the curve to better match experimental torque data. This may reflect inter-individual differences in joint morphology or muscle path constraints that are not fully captured by the generic model. Importantly, representing moment arm variation with a single scaling parameter appears sufficient to capture meaningful variability while maintaining model parsimony, supporting its inclusion in the hierarchical framework.

Previous studies have shown that EMG amplitudes do not follow a strictly linear relationship with muscle force during isometric contractions (Woods & Bigland-Ritchie, 1983). Our findings regarding the nonlinear EMG–activation exponents are consistent with this literature. Specifically, we obtained posterior estimates below 1.0 for both the elbow flexors and extensors (median ≈ 0.76 for flexors and ≈ 0.65 for extensors; Figure 5I–L). Because an exponent less than one implies a concave-downward relationship in the power function (Equation 6), increases in EMG amplitude yield diminishing gains in CE activation. Notably, the exponential formulation proposed by Lloyd & Besier (2003 (Lloyd & Besier, 2003)), within its reported parameter range, also yields a concave-down EMG–activation relationship, consistent with our power-function results. Together, these nonlinear formulations effectively capture the saturating relationship between muscle activation (or force) and EMG, particularly at higher target torque levels.

Compared with the EMG-driven model with the point-estimate method (Zhang et al., 2024) or MRI-based personalization methods, our Bayesian framework offers several advantages. Bayesian inference has long been recognized as a framework for modeling uncertainty in motor control, where variability in sensory input and motor output necessitates probabilistic reasoning (Kording & Wolpert, 2006). Our approach applies these principles to musculoskeletal modeling, enabling us to quantify uncertainty in parameter estimates and provide a richer understanding of the biomechanical characteristics of our participants. A central advantage of a Bayesian framework is the ability to incorporate prior knowledge into the parameter estimation process. The Bayesian approach achieved substantially better performance than the point-estimate method, particularly when only a limited number of trials from the test subjects were available. Here, we used prior distributions that were physiologically plausible and weakly informative, meaning they provided gentle regularization without strongly constraining the parameter values. Weakly informative priors incorporate broad biological knowledge—such as the expected scale of muscle forces or tendon stiffness—while remaining wide enough that the data, rather than the priors, primarily determine the posterior estimates. When we systematically varied the prior scale, the resulting posterior estimates remained highly stable. Across all prior settings, the percentage changes in parameter medians were generally within 5%, and only three individual-level parameters exhibited changes greater than 10% relative to the baseline model. This indicates that even when the priors were substantially tightened or made more diffuse, the inferred parameter values changed only minimally. Taken together, these results demonstrate that both the group-level and individual-level inferences are robust to prior specification, and that the primary conclusions reflect information contained in the data rather than sensitivity to the choice of weakly informative priors. In future studies, these priors could be further refined using empirical data from large musculoskeletal datasets or other MRI-based studies. For example, population-level distributions of tendon slack length or EMG nonlinearity relationships could inform more accurate prior means and variances.

Additionally, the hierarchical structure enables partial pooling across subjects, thereby improving estimates of muscle force relative to a non-hierarchical approach while preserving individual specificity. This approach has been shown to enhance inference in repeated-measures designs and in clinical studies with substantial inter-individual variability (Schweighofer et al., 2023; Veenman et al., 2024). In contrast to models that treat each individual entirely independently (no pooling) or assume all individuals share the same parameters (complete pooling), partial pooling allows individual parameter estimates to be informed by the group-level distribution. This approach stabilizes estimates for participants with noisier data while still preserving meaningful inter-individual differences. For example, in our model, subject-specific parameters, such as the Flexor Scale and Extensor Scale, were drawn from shared group-level distributions, enabling robust inference even when data quality varied across participants. This structure is particularly valuable in biomechanics, where physiological variability exists but is often constrained by shared anatomical and functional principles. Lastly, EMG-driven models have been widely used to estimate muscle forces, but they often require extensive calibration and are limited by the accessibility of deep muscles and the reliability of surface EMG signals (Ao et al., 2022; Hambly et al., 2025). Our Bayesian framework addresses these limitations by explicitly modeling uncertainty and inferring parameters, even in the presence of noisy or incomplete EMG data.

The EMG–torque relationship provides insight into motor-unit recruitment and muscle-activation strategies. Numerous studies have examined the relationship between surface EMG amplitude and joint torque (Ahamed et al., 2014; Clancy et al., 2006; Paquin & Power, 2018) and both linear and nonlinear relationships have been reported (Clancy et al., 2002; Woods & Bigland-Ritchie, 1983). Studies have shown that ignoring EMG coactivations can significantly influence the EMG-torque relationship (Brown & McGill, 2008); therefore, we included surface EMG from both the agonist and antagonist muscles in the analysis. However, this reliance on surface EMG introduces an inherent limitation: surface EMG cannot perfectly isolate deep or overlapping muscle activity, and residual crosstalk or imperfect representation of true neuromuscular activation patterns may still influence the inferred parameter estimates.

Despite its strengths, the current approach has limitations. First, EMG normalization and signal quality remain challenges, particularly for deep muscles or across sessions. Here, we focused on an isometric task, and as a result, our approach does not incorporate force-velocity dynamics, leaving the generalizability to dynamic tasks an open question. Second, we used lumped parameters to scale the tendon slack length and muscle moment arms for each of the five muscles in the model. This assumes that changes in these values would occur to the same extent within a participant, yet it is possible that there is variance among muscles within a subject in muscle moment arm profiles and tendon slack lengths that would not be captured by this approach. However, additional data, including imaging data, would be required to understand these relationships. Future work could extend this framework to dynamic movements, where time-varying kinematics and muscle forces introduce additional complexity.

Incorporating additional data modalities (e.g., ultrasound imaging to estimate muscle fascicle lengths) could improve model fidelity and expand the ability to perform these analyses outside of lab settings (Li & Tong, 2016). Direct measurements of fascicle length would provide information about muscle architecture and contractile state that surface EMG alone cannot capture. Because fascicle length influences force-length behavior, tendon stretch, and the mechanical moment transmitted to the joint, ultrasound measurements could greatly reduce uncertainty in key model parameters and improve the physiological interpretability of muscle–tendon dynamics in our framework.

## 5. Conclusion

This study presents a hierarchical Bayesian framework for estimating individualized muscle model parameters from EMG and torque data during isometric elbow tasks. By modeling both subject-specific and group-level variability, the approach enables robust inference of physiologically meaningful parameters while quantifying uncertainty in each estimate. Validation on synthetic data supports the model’s accuracy and convergence, and application to experimental data revealed substantial inter-individual differences in muscle strength and EMG-activation characteristics. Compared with traditional modeling approaches, this framework provides a scalable, interpretable method for personalizing musculoskeletal models, eliminating the need for imaging or invasive measurements. Future work should leverage Bayesian musculoskeletal modeling approaches to investigate clinical populations, such as people post-stroke, where neuromuscular impairments alter muscle coordination and force generation. In such cases, Bayesian inference could help quantify the extent of impairment and guide personalized rehabilitation strategies. As Bayesian methods continue to evolve, their integration into neuromechanical modeling holds promise for advancing both basic research and clinical applications in movement science.

## List of Abbreviations

MRI: magnetic resonance image
EMG: electromyography
MCMC: Markov Chain Monte Carlo
MVF: maximum voluntary force
DC: direct current
kHz: kilohertz
CE: Contractile Element
SEE: Series Elastic Element
F: Force
A: Activation
ES: Extensor Scale
FS: Flexor Scale
F-L: Force-Length
L_slack_: SEE slack length
*λ*: Force Scaling parameter that is applied to either the Extensors or Flexors
*Λ*: Anthropometry scale factor that adjusts lengths in the model based on participants’ height
W: width (referring to the width of the Force-Length curve)
P: Power (referring to the power term that dictates the non-linearity of the EMG-force relationship)
*β*: Scaling factor for the tendon slack length (Tendon Slack Scale)
*ψ*: scaling factor for the moment arm (Wave Scale)
MA: Moment Arm
JAGS: Just Another Gibbs Sampler
WAIC: Watanabe-Akaike Information Criteria
MPE: mean percent error
MAE: mean absolute error
RMSE: root-mean-squared error
R: a discrete parameter that is used to indicate the dominant or non-dominant arm
*ρ*: regression coefficient
HMC: Hamiltonian Monte Carlo
NUTS: No-U-Turn Samplers

## Declarations

### Ethics approval and consent to participate

- Experiments were performed with informed consent at the University of Southern California, with the approval of the local Institutional Review Board. Protocol ## Study ID: HS-05-00134

### Consent for publication

- All subjects gave their permission through written informed consent to publish individual data.

### Availability of data and materials

- All data are available in the main text or in supplementary materials. Code to reproduce the results found in this paper is available at https://github.com/russelljohnson95/bayesian_neuromechanics_25/tree/main

### Authors’ contributions

- Conceptualization: R.J., N.S., J.F.
- Designed experiments: Y.D., N.S., V.B.
- Conducted experiments: Y.D.
- Analyzed Data: R.J., Y.Y., Y.D., D.Y.
- Performed MCMC: R.J., Y.Y
- Interpreted Data: R.J., Y.Y., N.S., J.F.
- Prepared Initial Manuscript: R.J, Y.Y, N.S.
- All authors read and approved of the final manuscript.

## Acknowledgements

Not applicable

