## Supplementary Materials for "Estimating Muscle Parameters via Hierarchical Bayesian Neuromechanics"

**Supplement 1 Prior Sensitivity Tests**

***Methods***:

Prior sensitivity was assessed by rescaling the priors. For priors with a normal distribution, the prior standard deviation was multiplied by factors between 0.1 and 10 (10 log-spaced values). For priors with a gamma distribution, both the shape and scale parameters were multiplied by the same set of log-spaced scaling factors (0.1 to 10).

The hierarchical Bayesian model was then fitted under each prior specification, and the posterior medians of group-level and individual-level parameters are reported as percentage changes relative to those in the main manuscript.

***Results:***
The percentage changes of the parameter posterior medians comparing to the baseline are listed in Table 1 (group level parameters), and Table 2 (individual level parameters).

**Table S1.1.** Prior Sensitivity Test Results for the Group-Level Parameters

| Group-level parameters | Min and Max of the posterior median from sensitivity test | Percent change compared to the priors used in the manuscripts |
|---|---|---|
| $\mu_{ES}$ | [1.9,2.0] | [-2.3%,2.2%] |
| $\mu_{FS}$ | [2.1,2.2] | [-3.2%,0.9%] |
| $\mu_{\psi}$ | [0.86,0.87] | [-0.8%,0.1%] |
| $\mu_{TSL}$ | [1.1,1.2] | [-0.1%,0.1%] |
| $\mu_{\mathrm{EE}}$ | [0.64,0.66] | [-0.5%,1.2%] |
| $\mu_{\mathrm{FE}}$ | [0.76,0.77] | [-0.3%,0.1%] |
| $\sigma_{ES}$ | [0.63,0.70] | [-1.9%,17.7%] |
| $\sigma_{FS}$ | [0.89,0.94] | [-2.0%,9.0%] |
| $\sigma_{\psi}$ | [0.036,0.075] | [-26.0%,71.6%] |
| $\sigma_{TSL}$ | [0.030,0.035] | [-28.9%,3.8%] |
| $\sigma_{\mathrm{EE}}$ | [0.032,0.069] | [-14.3%,75.6%] |
| $\sigma_{\mathrm{FE}}$ | [0.032,0.054] | [-10.2%,62.2%] |

**Table S1.2**. Prior sensitivity test results for the individual-level parameters (Percentage Change Comparing to the posteriors reported in the main manuscript). Bolded entries indicate where changes to the prior resulted in estimates that were greater than 10% different from the nominal priors.

| Subject | FlexScale | ExtScale | WaveScale | TSL | Flexor Exponent | Extensor Exponent |
|---|---|---|---|---|---|---|
| S1 | [-0.5%,0.2%] | [-0.6%,8.0%] | [-1.3%,0.4%] | [-0.1%,0.1%] | [-0.3%,0.8%] | [-0.6%,5.7%] |
| S2 | [-1.1%,4.3%] | [-0.5%,3.9%] | **[-11.7%,2.4%]** | [-0.9%,0.1%] | [-0.5%,1.4%] | [-1.0%,5.9%] |
| S3 | [-0.6%,5.0%] | [-0.7,5.8%] | [-0.5%,2.6%] | [-0.2%,0.1%] | [-0.8%,6.5%] | [-0.8%,7.1%] |
| S4 | [-0.8%,0.0%] | [-3.0%,0.5%] | [-1.5%,0.4%] | [-0.1%,0.1%] | [-4.6%,0.2%] | [-6.2%,0.5%] |
| S5 | [-0.6%,2.0%] | [-2.7%,-0.1%] | [-0.5%,2.2%] | [-0.1%,0.1%] | [-0.8%,2.2%] | [-2.0%,-0.3%] |
| S6 | [-1.3%,0.2%] | [-2.1%,1.1%] | [-2.3%,0.5%] | [-0.1%,0.1%] | [-1.6%,0.1%] | [-1.9%,0.6%] |
| S7 | [-3.9%,0.2%] | [-1.3%,0.1%] | [-3.5%,0.4%] | [-0.2%,0.1%] | [-4.0%,0.3%] | [-1.8%,0.2%] |
| S8 | [-0.8%,0.9%] | [-0.9%,0.3%] | [-0.7%,2.3%] | [-0.2%,0.1%] | [-1.0%,0.9%] | [-0.9%,-0.3%] |
| S9 | [-0.7%,-0.1%] | [-1.1%,0.1%] | [-0.9%,0.2%] | [-0.1%,0.1%] | [-1.4%,0.1%] | [-1.5%,0.1%] |
| S10 | [-1.0%,2.9%] | [-0.2%,1.2%] | [-0.6%,0.2%] | [-0.1%,0.1%] | [-0.7%,2.2%] | [-0.6%,2.4%] |
| S11 | [-0.1%,1.0%] | [-0.3%,1.6%] | [-0.4%,1.1%] | [-0.1%,0.1%] | [-0.3%,1.9%] | [-0.4%,2.5%] |
| S12 | [-3.9%,0.7%] | [-1.5%,0.1%] | [-0.5%,0.8%] | [-0.1%,0.1%] | [-5.7%,0.4%] | [-5.8%,0.2%] |
| S13 | [-2.8%,0.2%] | [-1.3%,0.2%] | [-0.5%,1.5%] | [-0.2%,0.1%] | [-2.3%,-0.1%] | [-2.5%,0.4%] |
| S14 | [-3.3%,0.3%] | **[-2.0%,21.0%]** | [-2.7%,0.8%] | [-0.1%,0.1%] | [-0.7%,0.2%] | **[-1.7%,17.7%]** |

***Interpretation:***

Table 1.1 shows that the percentage change for all group-level means under different priors is within 5% of the baseline, indicating that the posterior group-level parameter medians are not sensitive to the choice of priors. For the group-level standard deviations, the percentage changes for the Force Scales and Tendon Slack Scale are below 20%. Although the percentage changes for Wave Scale and the two Exponents exceed 50%, their absolute differences across different prior conditions are small (all below 0.05).

Table 1.2 reports the percentage change in the posterior median of each individual-level parameter for each participant. In general, these percentage changes are within 5% (with only 3 individual posterior parameters having more than 10% change compared to the baseline), suggesting that the individual-level parameters are also robust to different prior specifications.

**Supplement 2 --- Sensitivity test on co-activation**

***Methods***:
Coactivation sensitivity was assessed by setting the EMG from the antagonist muscles to be 0, and then the hierarchical Bayesian model was fitted with the new EMG input, and the posterior medians of group-level and individual-level parameters are reported as percentage changes relative to those in the main manuscript.

***Results:***
The percentage changes of the parameter posterior medians compared to the baseline are listed in Table 2.1 (group level parameters), and Table 2.2 (individual level parameters).

**Table S2.1** Sensitivity test on co-activation for the Group-Level Parameters

| **Group-level parameters** | **Posterior Median (Original Model)** | **Posterior Median and percent change (Model without co-activation)** |
|---|---|---|
| $\mu_{ES}$ | 1.92 | 1.08 (-43.8%) |
| $\mu_{FS}$ | 2.16 | 1.29 (-40.3%) |
| $\mu_{\psi}$ | 0.87 | 0.95 (+9.2%) |
| $\mu_{TSL}$ | 1.18 | 1.17 (-0.80%) |
| $\mu_{\mathrm{EE}}$ | 0.65 | 0.52 (-20.0%) |
| $\mu_{\mathrm{FE}}$ | 0.76 | 0.67 (-11.8%) |
| $\sigma_{ES}$ | 0.61 | 0.54 (-11.5%) |
| $\sigma_{FS}$ | 0.87 | 0.63 (-27.6%) |
| $\sigma_{\psi}$ | 0.040 | 0.040 (0.0%) |
| $\sigma_{TSL}$ | 0.035 | 0.035 (0.0%) |
| $\sigma_{\mathrm{EE}}$ | 0.034 | 0.033 (-2.9%) |
| $\sigma_{\mathrm{FE}}$ | 0.033 | 0.034 (3.0%) |

**Table S2.2**. Coactivation sensitivity test results for the individual-level parameters (Percentage Change Comparing to the posteriors reported in the main manuscript).

| **Subject** | **FlexScale** | **ExtScale** | **WaveScale** | **TSL** | **Flexor Exponent** | **Extensor Exponent** |
|---|---|---|---|---|---|---|
| S1 | 2.14 (-20.1%) | 2.32 (-30.5%) | 0.86(2.4%) | 1.18 (0%) | 0.60 (-17.8%) | 0.57 (-20.8%) |
| S2 | 2.35 (-41.0%) | 1.51 (-24.5%) | 0.83 (18.6%) | 1.20 (0.8%) | 0.69 (-14.8%) | 0.55 (-20.3%) |
| S3 | 1.24 (-38.3%) | 1.13 (-36.2%) | 1.01 (9.8%) | 1.16 (-1.7%) | 0.76 (-10.6%) | 0.58 (-14.7%) |
| S4 | 0.32 (-67.3%) | 0.51 (-55.7%) | 1.08 (27.1%) | 1.16 (-2.5%) | 0.71 (-6.6%) | 0.53 (-8.6%) |
| S5 | 0.74 (-47.9%) | 0.71 (-39.8%) | 1.00 (6.38%) | 1.19 (-1.7%) | 0.76 (-7.3%) | 0.50 (-10.7%) |
| S6 | 1.06 (-57.8%) | 1.13 (-36.5%) | 0.95 (17.3%) | 1.13 (-3.4%) | 0.69 (-10.4%) | 0.50 (-18.0%) |
| S7 | 1.33 (-27.3%) | 0.97 (-49.2%) | 0.88 (7.3%) | 1.17 (-0.8%) | 0.63 (-10.0%) | 0.50 (-25.3%) |
| S8 | 1.27 (-39.8%) | 1.04 (-60.0%) | 0.95 (-2.1%) | 1.20 (0.8%) | 0.66 (-16.5%) | 0.44 (-26.7%) |
| S9 | 1.02 (-39.6%) | 1.03 (-50.7%) | 0.91 (9.6%) | 1.12 (-1.8%) | 0.63 (-14.9%) | 0.47 (-25.4%) |
| S10 | 1.91 (-44.5%) | 0.75 (-59.2%) | 0.89 (7.2%) | 1.21 (-1.6%) | 0.72 (-8.9%) | 0.50 (-23.1%) |
| S11 | 1.68 (-37.3%) | 1.37 (-32.5%) | 0.98 (6.5%) | 1.19 (-0.8%) | 0.66 (-14.3%) | 0.52 (-23.5%) |
| S12 | 1.43 (-27.4%) | 1.32 (-27.9%) | 1.00 (6.4%) | 1.17 (-0.8%) | 0.58 (-13.4%) | 0.47 (-20.3%) |
| S13 | 1.92 (-31.2%) | 1.81 (-27.3%) | 1.01 (7.4%) | 1.18 (0.9%) | 0.62 (-13.9%) | 0.51 (-19.0%) |
| S14 | 1.19 (-24.7%) | 1.05 (-16.7%) | 0.89 (9.9%) | 1.18 (0%) | 0.61(-15.3%) | 0.59 (-20.3%) |

**Supplement 3:**

***Methods***: To assess independence of sampling across parameters from the synthetic data set, we computed pairwise Pearson's correlation coefficients among all posterior samples (not medians). Low correlations indicate that the sampler explores each parameter's solution space independently, which reflects good mixing and minimal interdependence in the sampling process.

***Results:*** Figure S3.1 shows the correlation matrix of posterior samples for all six parameters. All correlations are near zero, with the highest value below 0.35. This pattern demonstrates that the sampling algorithm moves independently through each parameter's solution space, with little coupling between parameters.

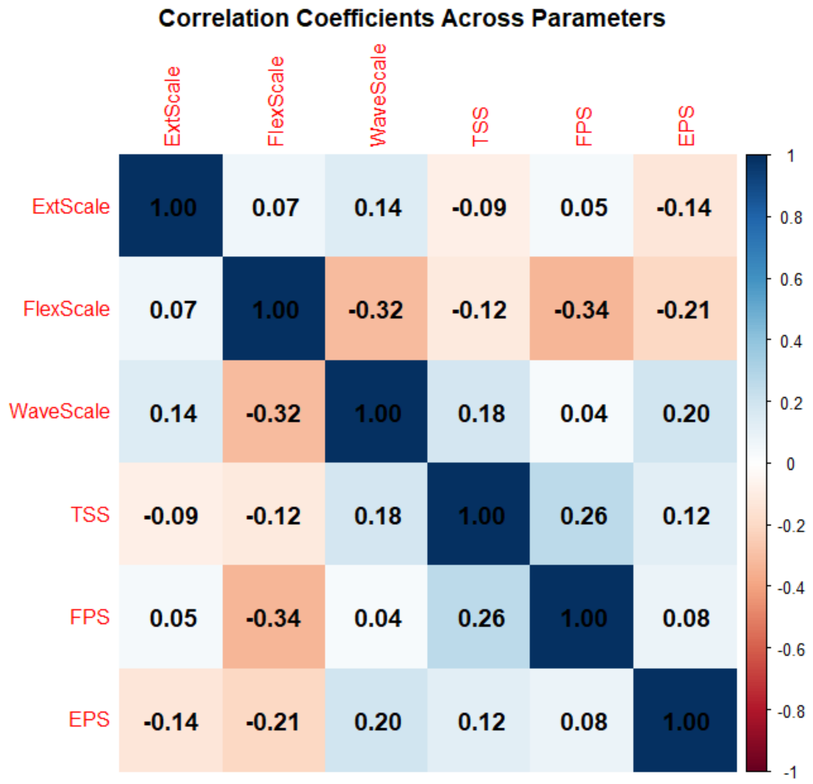


**Figure S3.1**: Correlation matrix of posterior samples for six model parameters. Low correlations indicate that the sampling algorithm explores each parameter's solution space independently, demonstrating good mixing and minimal interdependence among parameters.

***Interpretation:***

The low correlations among posterior samples confirm that the RJAGS sampler efficiently explores the joint posterior distribution. Each parameter is sampled largely independently, reducing the risk of biased estimates or poor mixing. High correlations would have indicated strong interdependence, potentially compromising identifiability and convergence. These results support the robustness of the sampling method and the reliability of the posterior estimates.

**Supplement 4: EMG-Torque Relationship Plot with strong co-activation**

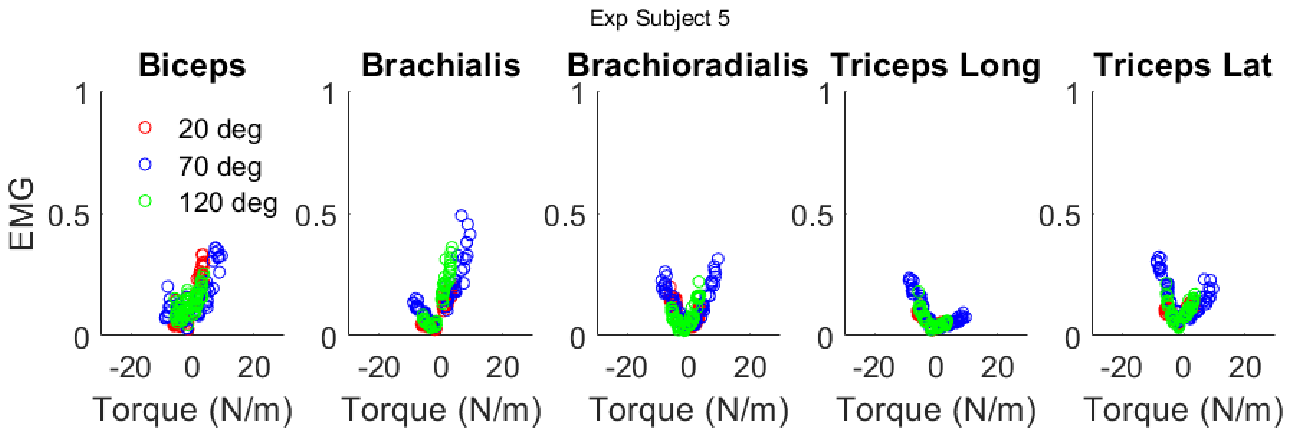


**Figure S1.1:** Surface EMG – Torque relationship plot from one subject who exhibited stronger muscle co-activation for both flexion and extension tasks. The y-axis is the average amplitude of the normalized surface EMG (normalized by MVC) within each trial, and the x-axis is the measured net elbow joint torque (Extension torques are plotted as negative values) for each trial.

### Supplement 5 Torque Recovery

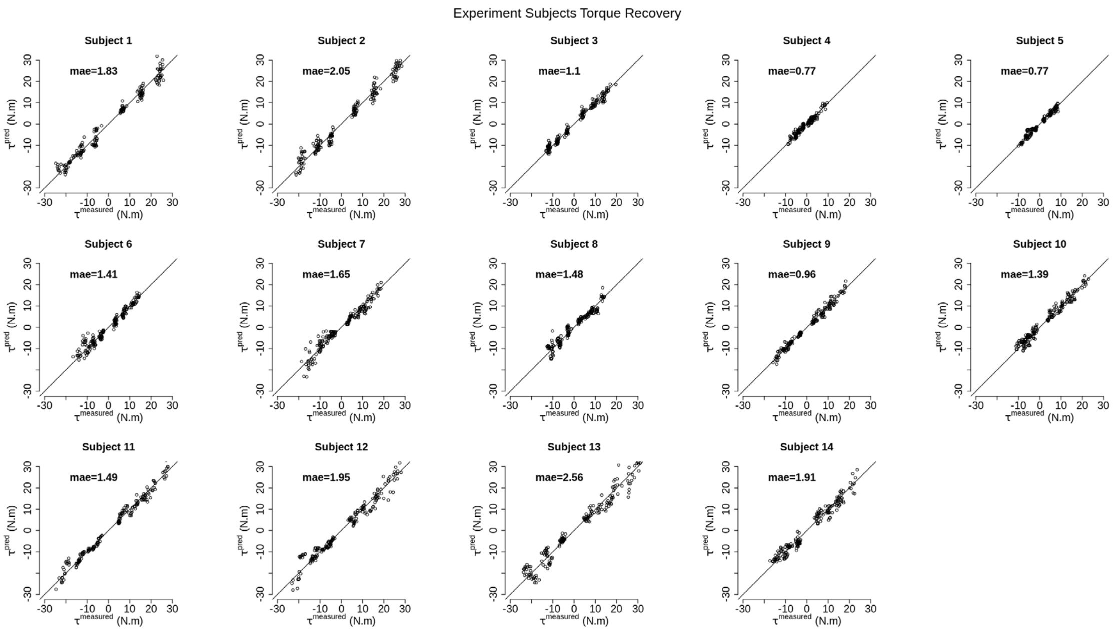


**Figure S5.1:** The x-axis is the measured torque from the experiment dataset. The y-axis is the predicted torque that is recovered via the Bayesian inference model. Each dot represents a single trial from an experimental subject; dots lying along the line of identity suggest that the error is equal between the measured torque and the torque recovered via Bayesian inference.

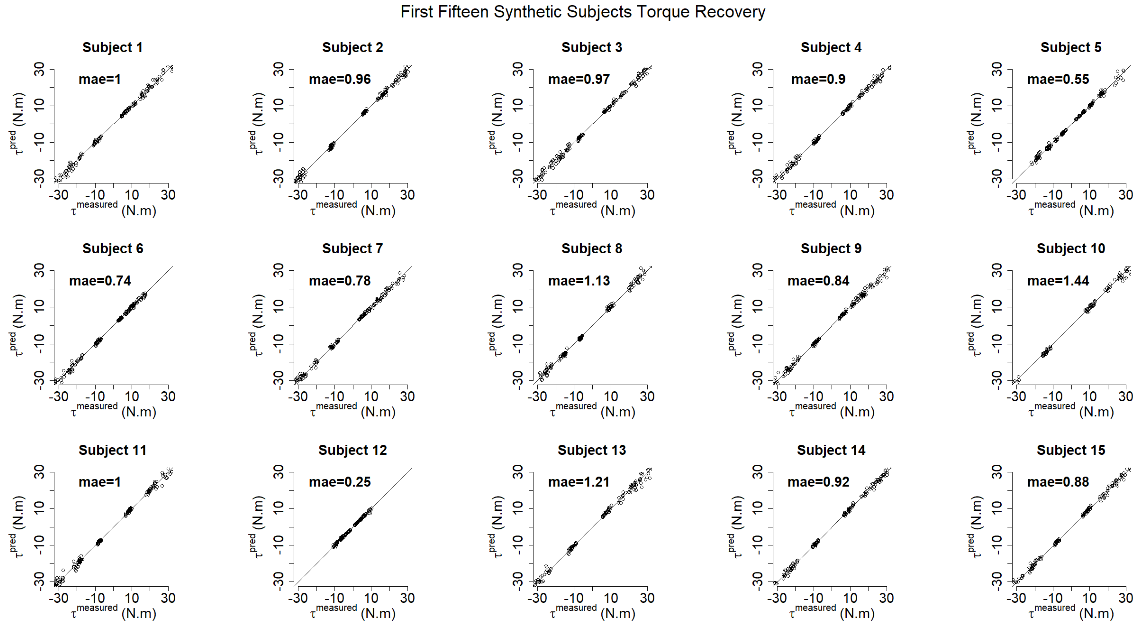


**Figure S5.2.** The x-axis is the measured torque from the synthetic dataset. The y-axis is the predicted torque that is recovered via the Bayesian inference model. Each dot represents a single trial from a synthetic participant; dots lying along the line of identity suggest that the error is equal between the measured torque and the torque recovered via Bayesian inference. Here, we show the first fifteen synthetic subjects as a representation of good torque recovery.

**Supplement 6: Exemplary Posterior Predictive Check**

**Methods:** Posterior predictive checks were performed to evaluate the adequacy of the hierarchical Bayesian model in capturing observed torque profiles. We generated predicted torque curves using posterior samples for two model configurations: (i) a reduced 4-parameter model and (ii) a full 6-parameter model. Predictions were compared against observed data for flexion and extension trials. Visual checks were used to assess whether the observed data fell within the 95% credible intervals of the posterior predictive distributions.

**Results:** Figures S5.1.1–S5.1.4 show posterior predictive checks for flexion and extension under the 4-parameter and 6-parameter models for subject 13, whose data has the largest fitting error according to Figure S2.1. Figures S5.2.1–S5.2.4 show posterior predictive checks for flexion and extension under the 4-parameter and 6-parameter models for subject 4, whose data has the smallest fitting error according to Figure S2.1.

For the 6-parameter model, predicted torque curves closely match observed data, with most observations falling within the 95% credible intervals. The 4-parameter model exhibits larger deviations, particularly in extension trials, indicating reduced flexibility in capturing subject-specific variability.

**Interpretation:** Posterior predictive checks confirm that the hierarchical Bayesian model provides accurate predictions of joint torque profiles. The 6-parameter model demonstrates a superior fit compared to the 4-parameter model, suggesting that the inclusion of additional parameters improves the representation of muscle-tendon dynamics. These results support the validity of the hierarchical Bayesian approach and highlight the importance of model complexity for capturing biomechanical variability.

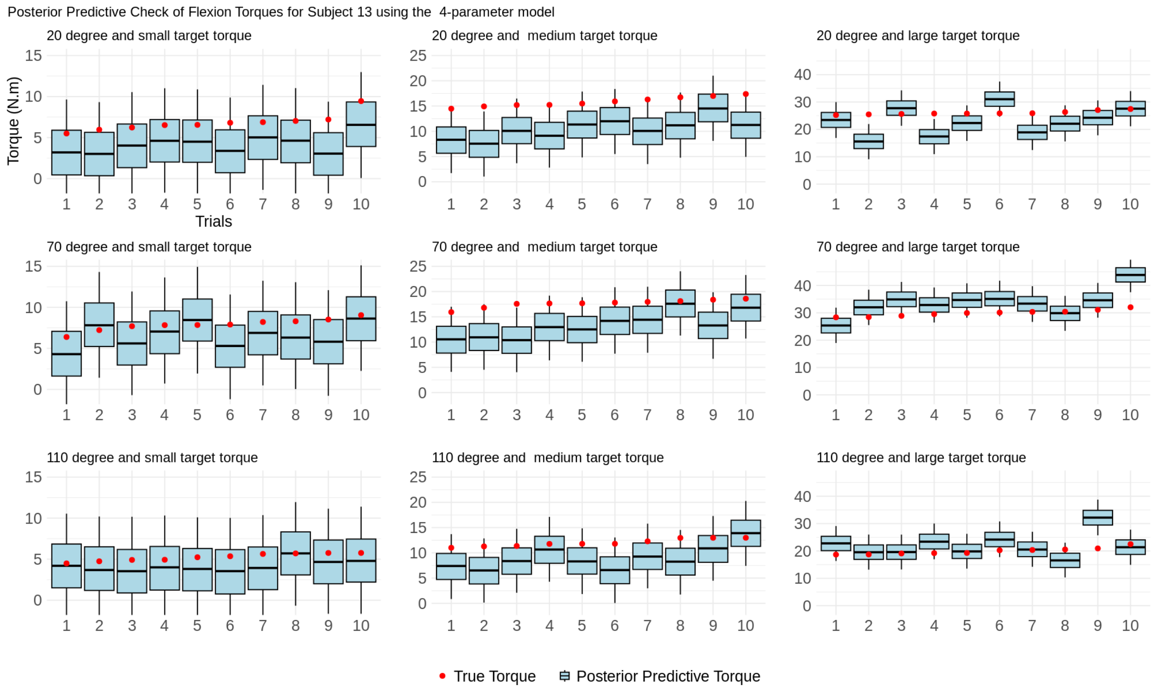


**Figure S6.1.1:** Posterior predictive check for Subject 13 using the 4-parameter model during flexion trials. Observed torque curves are compared against predictions with 95% credible intervals. The red dots are the true torques, the blue boxes are the 95% CI for the posterior predicted torques, and the black lines in the middle of the box are the median of the posterior predicted torques.

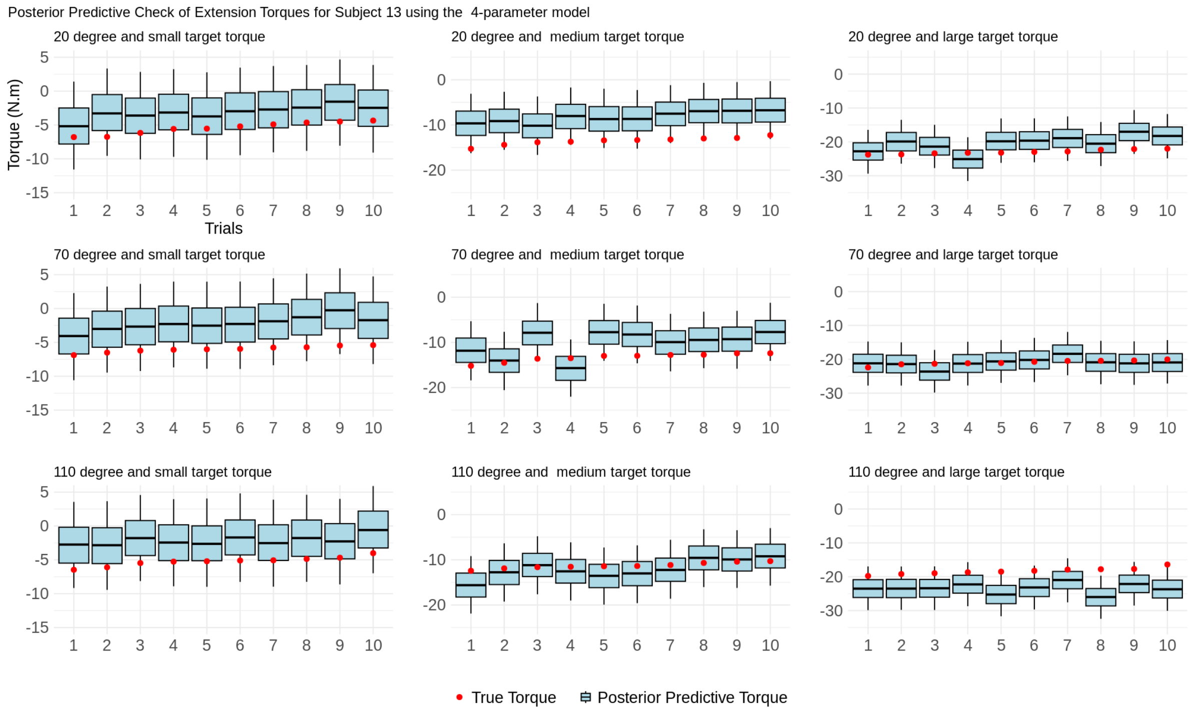


**Figure S6.1.2:** Posterior predictive check for Subject 13 using the 4-parameter model during extension trials. Deviations indicate reduced flexibility in capturing subject-specific variability.

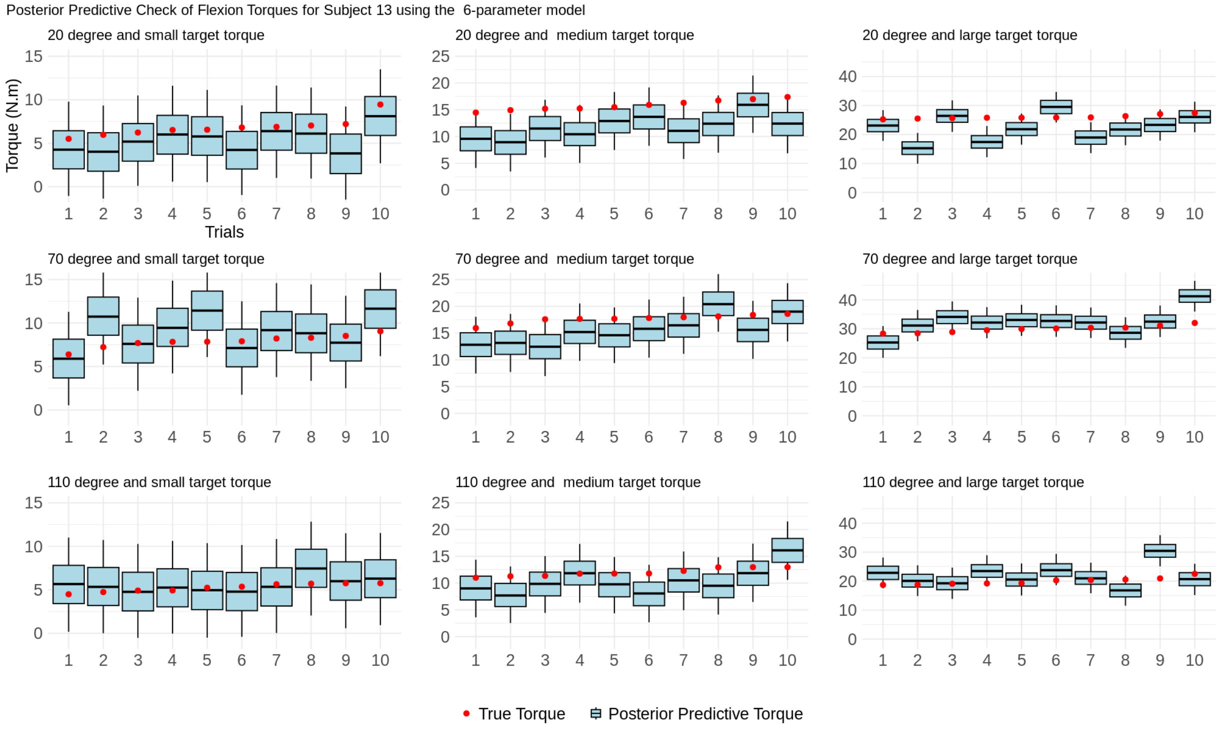


**Figure S6.1.3:** Posterior predictive check for Subject 13 using the 6-parameter model during flexion trials.

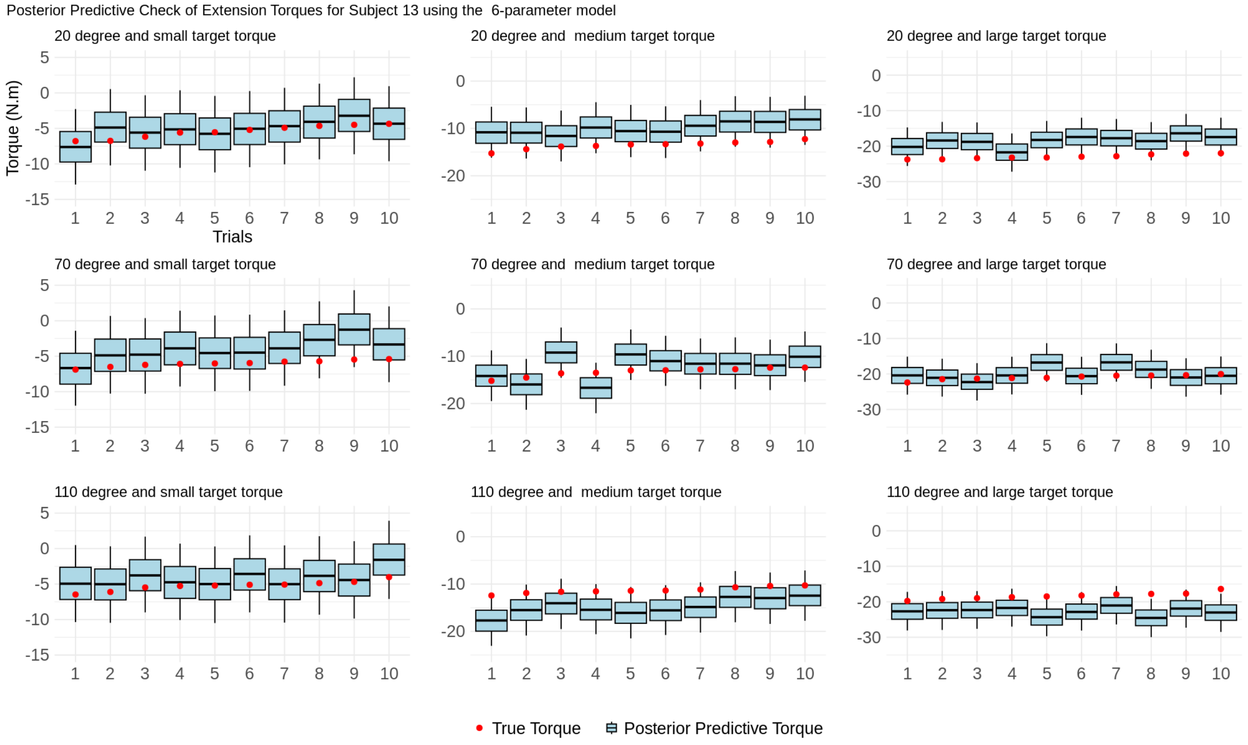


**Figure S6.1.4:** Posterior predictive check for Subject 13 using the 6-parameter model during extension trials.

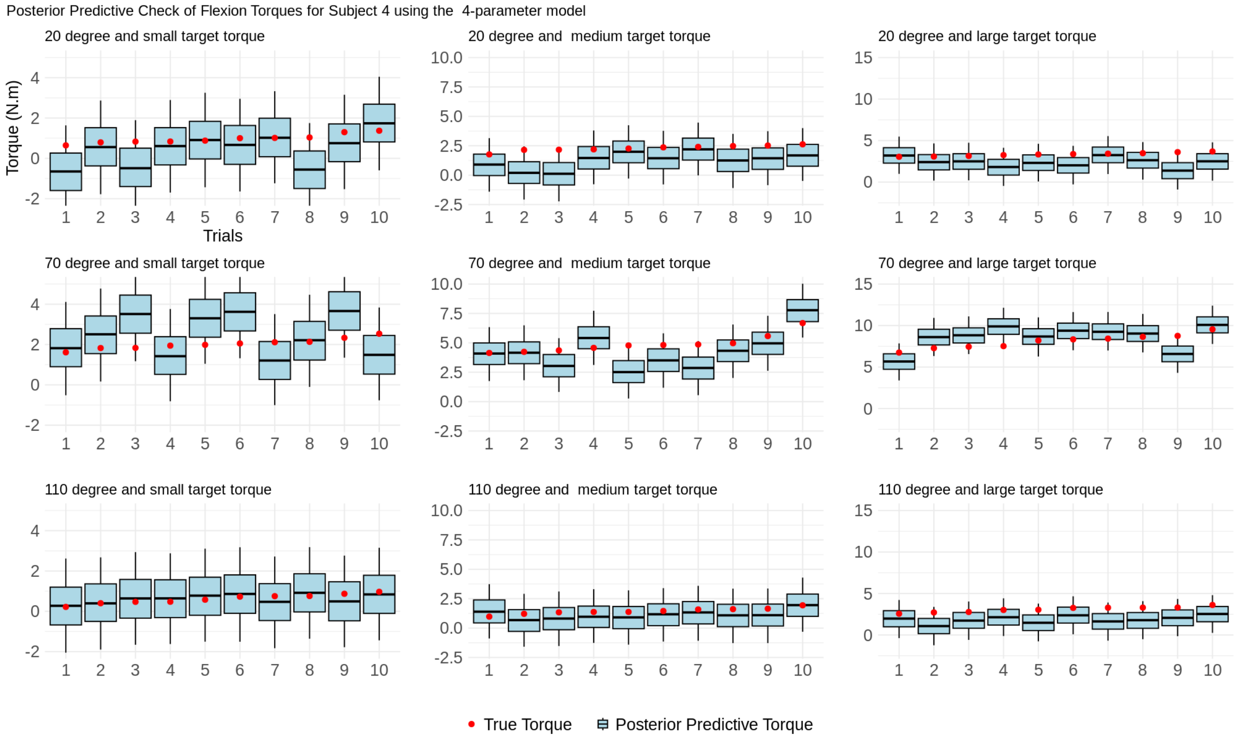


**Figure S6.2.1:** Posterior predictive check for Subject 4 using the 4-parameter model during flexion trials.

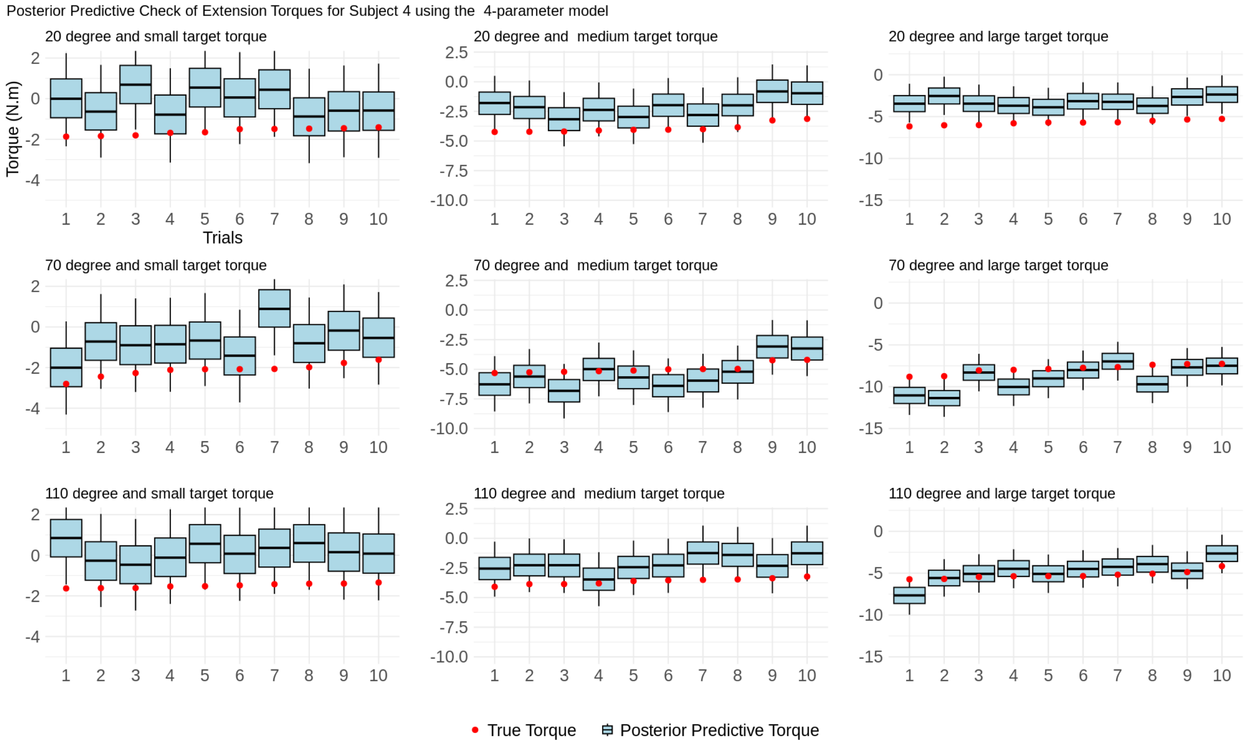


**Figure S6.2.2:** Posterior predictive check for Subject 4 using the 4-parameter model during extension trials.

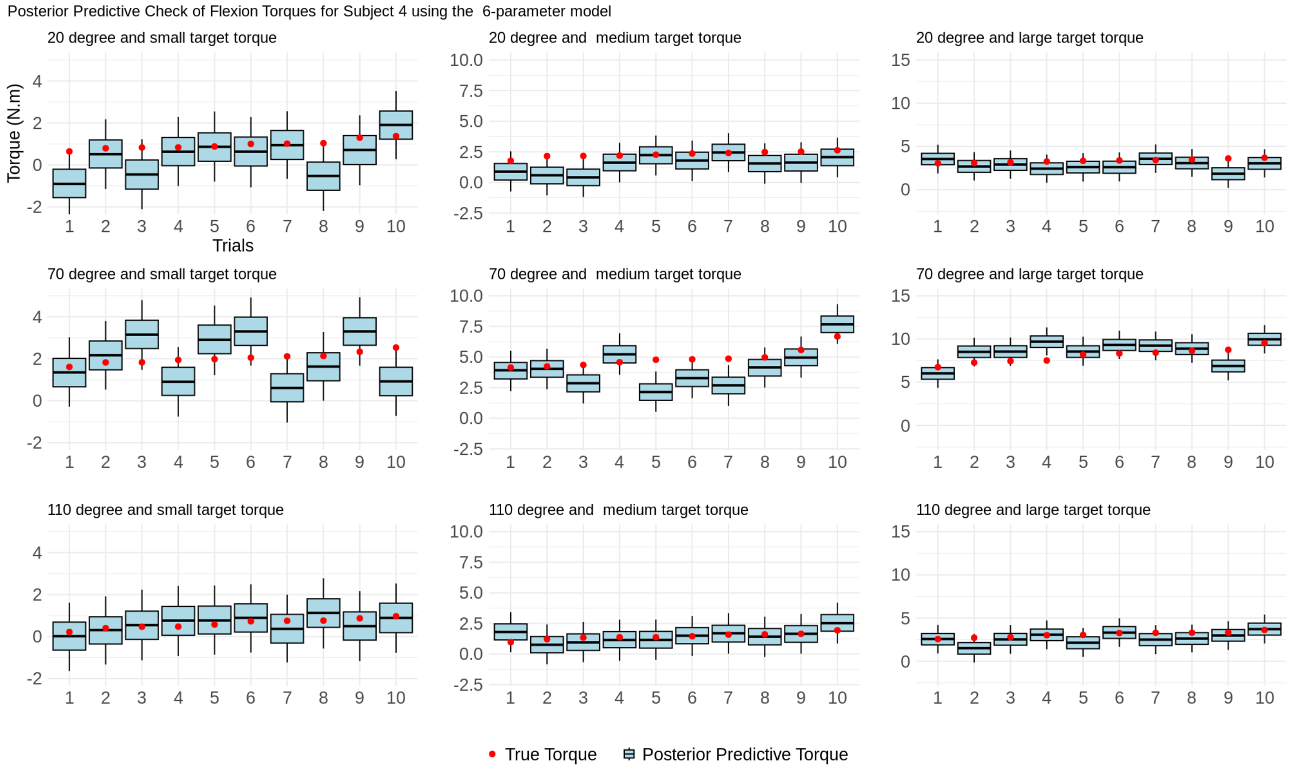


**Figure S6.2.3:** Posterior predictive check for Subject 4 using the 6-parameter model during flexion trials.

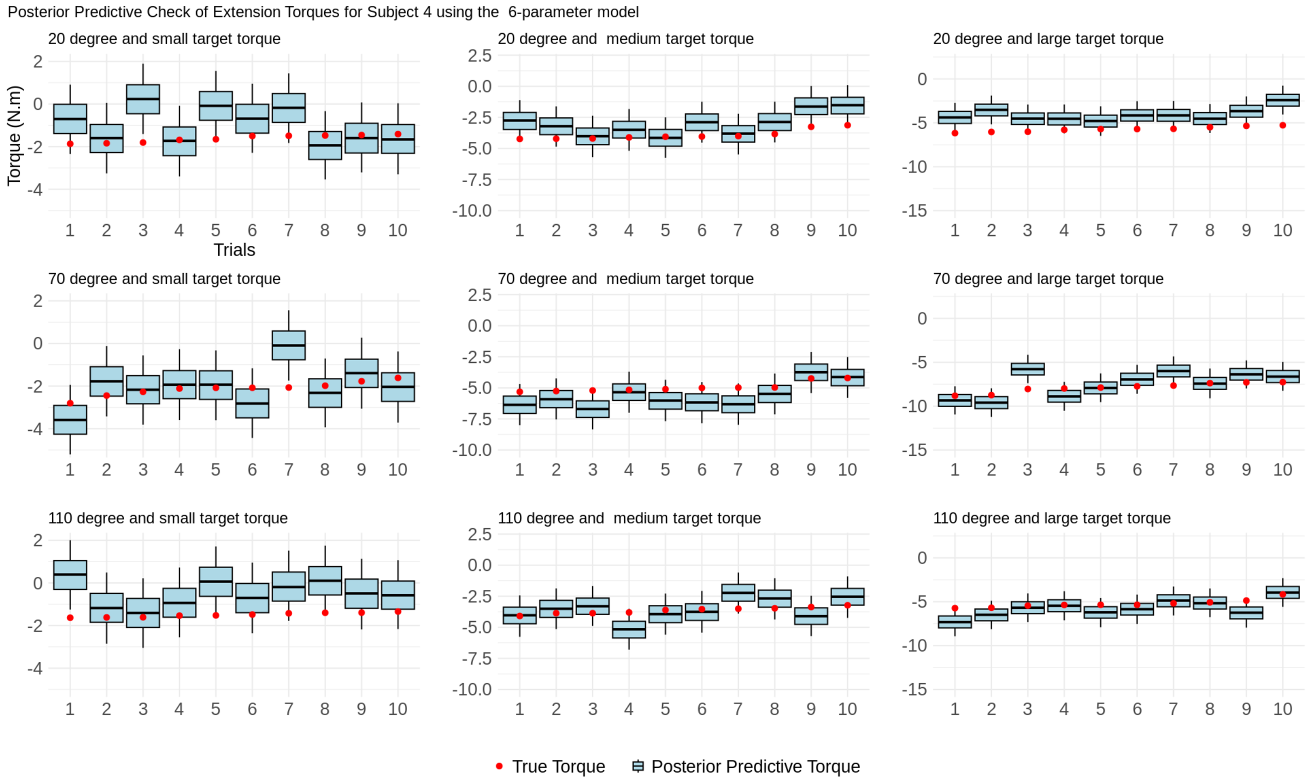


**Figure S6.2.4:** Posterior predictive check for Subject 4 using the 6-parameter model during extension trials.
